# Epigenetic priming of embryonic enhancer elements coordinates developmental gene networks

**DOI:** 10.1101/2024.09.09.611867

**Authors:** Christopher D Todd, Jannat Ijaz, Fereshteh Torabi, Oleksandr Dovgusha, Stephen Bevan, Olivia Cracknell, Tim Lohoff, Stephen Clark, Ricard Argelaguet, Juliette Pearce, Ioannis Kafetzopoulos, Alice Santambrogio, Jennifer Nichols, Ferdinand von Meyenn, Ufuk Guenesdogan, Stefan Schoenfelder, Wolf Reik

**Author notes:** co-corresponding author, Correspondence: CDT, SS and WR.

## Abstract

Embryonic development requires the accurate spatiotemporal execution of cell lineage-specific gene expression programs, which are controlled by transcriptional enhancers. Developmental enhancers adopt a primed chromatin state prior to their activation; however how this primed enhancer state is established, maintained, and how it affects the regulation of developmental gene networks remains poorly understood. Here, we use comparative multi-omic analyses of human and mouse early embryonic development to identify subsets of post-gastrulation lineage-specific enhancers which are epigenetically primed ahead of their activation, marked by the histone modification H3K4me1 within the epiblast. We show that epigenetic priming occurs at lineage-specific enhancers for all three germ layers, and that epigenetic priming of enhancers confers lineage-specific regulation of key developmental gene networks. Surprisingly in some cases, lineage-specific enhancers are epigenetically marked already in the zygote, weeks before their activation during lineage specification. Moreover, we outline a generalisable strategy to use naturally occurring human genetic variation to delineate important sequence determinants of primed enhancer function. Our findings identify an evolutionarily conserved program of enhancer priming and begin to dissect the temporal dynamics and mechanisms of its establishment and maintenance during early mammalian development.

## Introduction

Embryogenesis requires the generation of a complex array of different cell types with distinct transcriptional programs. Enhancers, *cis*-acting transcriptional regulatory elements (1), play an important role in the programmed establishment and maintenance of cell fates through binding of sequence-specific transcription factors (TFs) and regulating transcription of their target genes in a cell-type-specific manner (2). Previous studies have utilised genetic analysis within disease models to highlight the potential functional importance of enhancer elements, such as the zone of polarizing activity regulatory sequence (ZRS) enhancer for the *Shh* gene (3). Genetic editing and deletion of this enhancer element, resulting in polydactyly and syndactyly, has provided conclusive evidence of this enhancer’s importance in establishing the correct spatiotemporal regulation of developmental tissues (4). While whole genome sequencing has facilitated the investigation of genetic factors such as TF binding motif sequences in enhancer activity, large-scale profiling of epigenetic marks has started to reveal the role of specific chromatin modifications in regulating enhancer activity. For example, active enhancers promoting transcription of associated genes are associated with the presence of the histone mark H3K27ac (5–8).

Studies of developmental programs have shown that lineage-specific enhancers often transition through pre-active states ahead of their subsequent activation; this “primed” enhancer state is defined as enhancers marked with the neutral enhancer-associated histone modification H3K4me1 in the absence of the active enhancer mark H3K27ac (9,10). A distinct “poised” pre-active state is defined as enhancers marked with primed (H3K4me1) and repressive (H3K27me3) marks (7,9). These pre-active enhancer states are hypothesised to prepare the enhancer region for activation upon binding of lineage-specific TFs to facilitate cell-fate associated changes of transcriptional programs. This is seen in various developmental processes including *Drosophila* mesoderm formation, zebrafish embryogenesis, neural crest development, cardiac lineage formation, and lymphogenesis (11–15).

Recent functional genomics and multi-omics approaches have begun to unravel the epigenetic regulation of enhancer activity and its role in early mammalian development. For example, tracing the epigenetic dynamics of lineage-specific enhancer elements during mouse gastrulation has revealed priming by DNA hypomethylation and accessible chromatin at neuroectodermal enhancers, but not mesodermal or endodermal enhancers, within the epiblast (16). However, several pertinent questions remain surrounding enhancer priming and its role in mammalian embryonic development. These include the following: (i) How is enhancer priming established and maintained? (ii) How do lineage-specific enhancers undergo priming during human embryonic development? (iii) What are the temporal dynamics of enhancer priming? In this study, we address these questions through multi-omic analysis of *in vitro* and *in vivo* models of human and mouse early embryonic development to determine the dynamics of enhancer epigenetic priming, to investigate its role in regulating cell fate decisions, and to identify potential trans-acting factors that establish the primed state at key human developmental enhancers.

## Results

### Epigenetic priming of lineage-specific enhancers in human and mouse epiblast

To identify epigenetic states of enhancer elements within the epiblast of human embryonic development, we analysed publicly available epigenomic data collected from post-gastrulation stage tissues differentiated from human epiblast-like embryonic stem cells (hESCs) (17). Lineage-specific enhancer elements for hESC-derived mesendoderm (hME) and neural progenitor stem cells (hNPC) were called from a previous list of random-forest model generated lineage-enriched enhancers (17), which were then subset by those which displayed lineage-specific H3K27ac (active enhancer) signals. This selection resulted in groups of predicted enhancer elements (1957 hNPC-specific and 4344 hME-specific enhancers) that appear epigenetically active only upon differentiation into their respective post-gastrulation lineage (Figure 1A). As a control group, we selected hESC enhancers with H3K27ac signal exclusively within the hESC lineage (i.e. absent in the post-differentiation lineages), resulting in 3750 hESC-specific enhancer elements (Figure 1A). We next looked at additional histone modification signatures within these lineage-specific enhancer groups and characterised their activity state in hESCs, categorising them either as: Active (H3K27ac+/H3K4me1+/H3K27me3-), Poised (H3K27ac-/H3K4me1+/H3K27me3+), Primed (H3K27ac-/H3K4me1+/H3K27me3-), or Inactive (H3K27ac-/H3K4me1-/H3K27me3-) (Supp. Fig.1-3). We identified a substantial subset of hNPC-specific and hME-specific enhancers displaying a Primed epigenetic signature within hESCs (27.4% of hNPC-specific enhancers/ 20.3% of hME enhancers) (Figure 1A), which we hereafter refer to as epiblast Primed (ePrimed) enhancers. For comparative analysis we generated a similarly sized subset of enhancers from the Inactive group which had the lowest H3K4me1 signal within the hESCs (Figure 1A), which we hereafter refer to as epiblast Non-primed (eNon-primed). We note that while these eNon-primed enhancers are not epigenetically primed within the epiblast, they may still undergo priming ahead of their activation at a time point not captured within this dataset.

**Figure 1.**
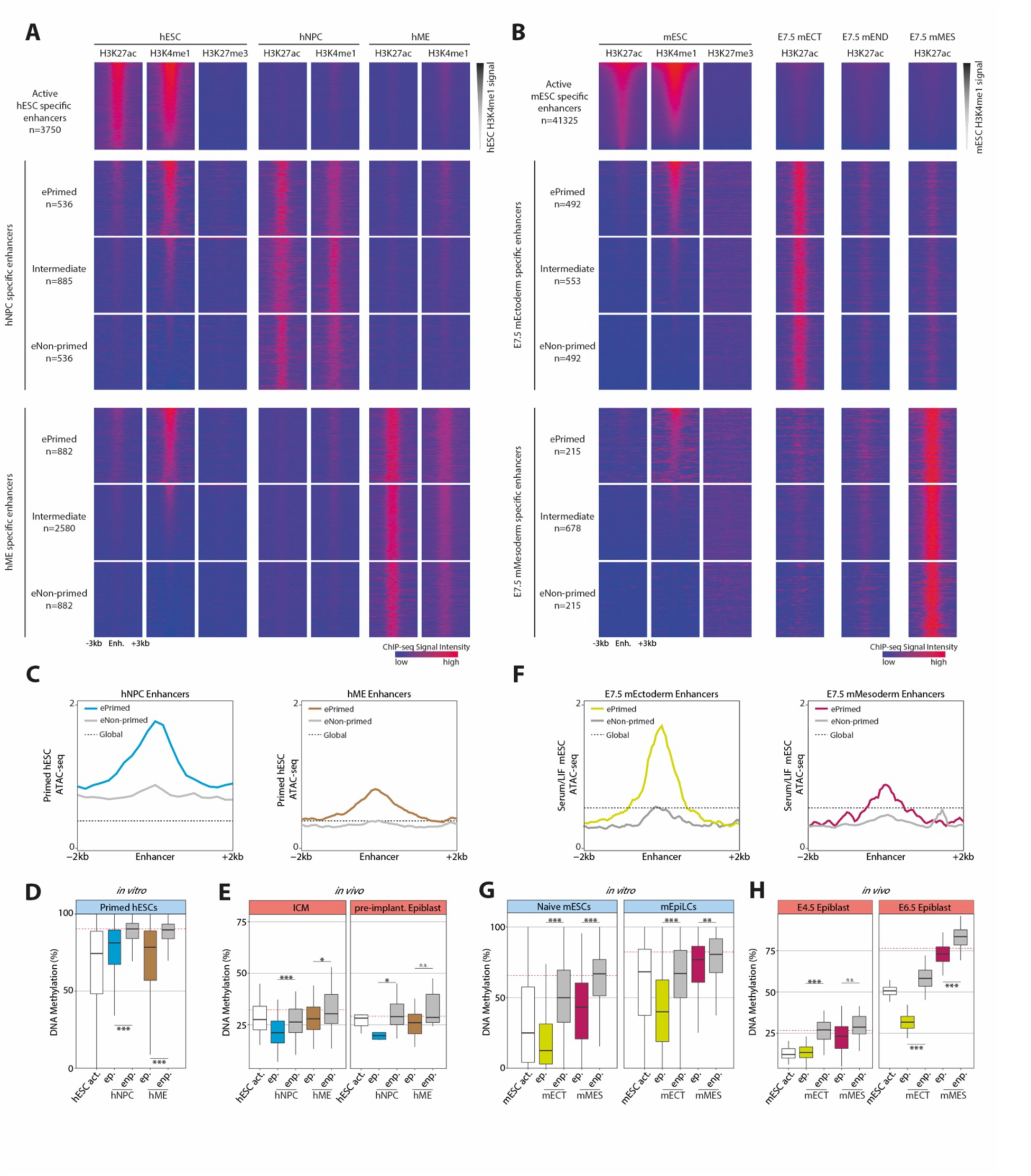
**(A)** Heatmaps of ChIP-seq H3K27ac, H3K4me1 and H3K27me3 data within hESC, hNPC and hME cells in culture at enhancer elements with lineage-specific active states in hESC, hNPC or hME cells. Heatmaps are ordered by H3K4me1 signal at the enhancer within hESCs. **(B)** Heatmaps of ChIP-seq H3K27ac, H3K4me1 and H3K27me3 data within mESCs, and of H3K27ac data within *in vivo* dissections of mECT, mEND, and mMES tissues for E7.5 embryos. Shown here are enhancer elements with lineage-specific active states in mESC cells, E7.5 mECT or E7.5 mMES tissues. Heatmaps are ordered by H3K4me1 signal at the enhancer within mESCs. **(A&B)** Enhancer groups called as ePrimed (high H3K4me1, low H3K27ac), eNon-primed (low H3K4me1, low H3K27ac), or Intermediate (intermediate H3K4me1) are displayed. **(C)** ATAC-seq profile for hNPC and hME called enhancer groups within *in vitro* primed hESCs. Running averages in 50-bp windows around the centre of the enhancer (2 kb upstream and downstream) are shown alongside the averaged global signal (black dashed line). **(D&E)** DNA methylation levels over 500bp core of active hESC enhancers (white) and hNPC and hME called enhancer sub-groups: ePrimed (ep.) and eNon-primed (enp.) **(D)** by WGBS within *in vitro* primed hESC and **(E)** by scNMT-seq of *in vivo* human preimplantation embryos. **(F)** ATAC-seq profile for E7.5 mEct and E7.5 mMES called enhancer groups within *in vitro* mESCs. Running averages in 50-bp windows around the centre of the enhancer (2 kb upstream and downstream) are shown alongside the averaged global signal (black dashed line). **(G&H)** DNA methylation levels over 500bp core of active mESC enhancers (white) and mECT and mMES called enhancer sub-groups: ePrimed (ep.) and eNon-primed (enp.) **(G)** by WGBS within *in vitro* naive mESC/mEpiLCs and **(H)** by scNMT-seq of *in vivo* mouse embryos at days E4.5 and E6.5. **(D-E,G-H)** Box plots show median levels and the first and third quartile, whiskers show 1.5x the interquartile range. Global methylation levels are displayed (red dashed line) along with results of ANOVA test between ePrimed and eNon-primed groups (p<0.001 ***, p<0.05 *).

DNA methylation and chromatin accessibility analysis of ePrimed enhancer subsets within hESCs identified additional associated signatures, including increased chromatin accessibility (Figure 1C), and decreased DNA methylation levels (Figure 1D) when compared to eNon-primed enhancers from the same lineage group. Notably, single-cell multi-omic analysis (scNMT-seq) of *in vivo* preimplantation human embryos shows similar DNA hypomethylation states at ePrimed enhancer sites (Figure 1E), suggesting that the epigenetic priming signature we observe within *in vitro* cultures recapitulates a regulatory program present within *in vivo* embryonic development. Interestingly, priming of lineage-specific enhancer elements was not observed to result in stronger activation of the enhancers upon differentiation, with little correlation between H3K4me1 signal in the hESC and the H3K27ac levels in the differentiated cell types (Supp. Figure 4A). However, hESC H3K4me1 levels did correlate with hESC H3K27ac levels, with ePrimed enhancers showing a weak enrichment of H3K27ac signal over eNon-primed enhancers (Supp. Figure 4B), albeit at significantly lower levels than those present at active enhancers. This could suggest that there is either mild or transient activation of ePrimed enhancers as part of establishment/maintenance of the Primed enhancer state.

To identify lineage-specific enhancer histone modification dynamics in mouse embryonic enhancers and compare them to epigenetic states of lineage-specific enhancer elements in the mouse embryonic epiblast, we used existing lineage-specific promoter-distal H3K27ac data from E7.5 ectoderm (mECT), endoderm (mEND), and mesoderm (mMES) (16), and then analysed the epigenetic signatures of these sites within epiblast-like mouse embryonic stem cells (mESCs) in naive culture conditions (Figure 1B, Supp. Fig.3C-D, Supp. Figure 5). Similar to our analysis of human enhancers, we identified a substantial subset of lineage-specific enhancers for all three germ layers that display a Primed epigenetic signature (H3K27ac-/H3K4me1+/H3K27me3-) in mESCs. Like the human ePrimed enhancers (Figure 1C-D), the mouse ePrimed enhancers display increased chromatin accessibility and decreased DNA methylation levels, when compared to global levels and eNon-Primed enhancers, both within *in vitro* culture and *in vivo* single-cell multi-omic datasets (Figure 1F-H, Supp. Figure 5). Additionally, we observed a similar weak enrichment of H3K27ac signal at H3K4me1 marked ePrimed enhancers over eNon-primed enhancers (Supp. Figure 4C-D). These results suggest that profiling of H3K4me1 is able to identify subsets of endoderm and mesoderm enhancers undergoing priming signatures which are otherwise obscured within prior class-wide chromatin accessibility and DNA methylation analyses (16). Consistent with previous findings [6], these priming signatures are more pronounced within ectoderm and ectoderm-derived neural lineage enhancers, especially within later epiblast tissues (Figure 1C-D,F,H), compared to those observed at endoderm/mesoderm lineage enhancers.

### Primed enhancers are associated with lineage-specific regulation of developmental gene networks

To identify the impact of epigenetic priming of lineage enhancers on associated gene networks, we attempted to assign enhancers to the promoters they likely control. In the absence of chromatin conformation data from the post-epiblast tissues we are investigating, we utilised a combination of genomic proximity models and Promoter Capture Hi-C data of mEpiLCs/hESCs to assign enhancers to promoters they likely interact with, as previous studies have shown that enhancers can interact with promoters prior to their activation (18,19). We then further subset these for genes found to be interacting with exclusively ePrimed or eNon-primed enhancer subgroups. To determine the functional role of gene networks associated with ePrimed enhancers, we performed a gene ontology enrichment analysis. This showed that ePrimed enhancers were enriched for associations with developmental gene networks (Figure 2A-B). Genes exclusively associated with ePrimed hNPC enhancers are enriched for ectoderm differentiation and various neural development relevant processes such as Hedgehog signalling pathway and postsynaptic density (Figure 2A). Conversely, eNon-primed hNPC exclusively associated genes were not found to be enriched for any specific biological processes. The ePrimed hME enhancer-associated genes are similarly enriched for tissue-relevant processes, with enrichments for endoderm differentiation genes along with signalling pathways and processes related to axon development, which are important for the mesoderm’s function in establishing the neural tube (20)(Figure 2B). Again, hME eNon-primed enhancers were not enriched for any specific biological process. Similarly in mouse, ePrimed mECT/mEND enhancer-associated genes were enriched for morphogenic and developmental processes whereas eNon-primed mECT/mEND enhancer-associated genes were not enriched for any specific processes (Supp. Figure 5A-B).

**Figure 2.**
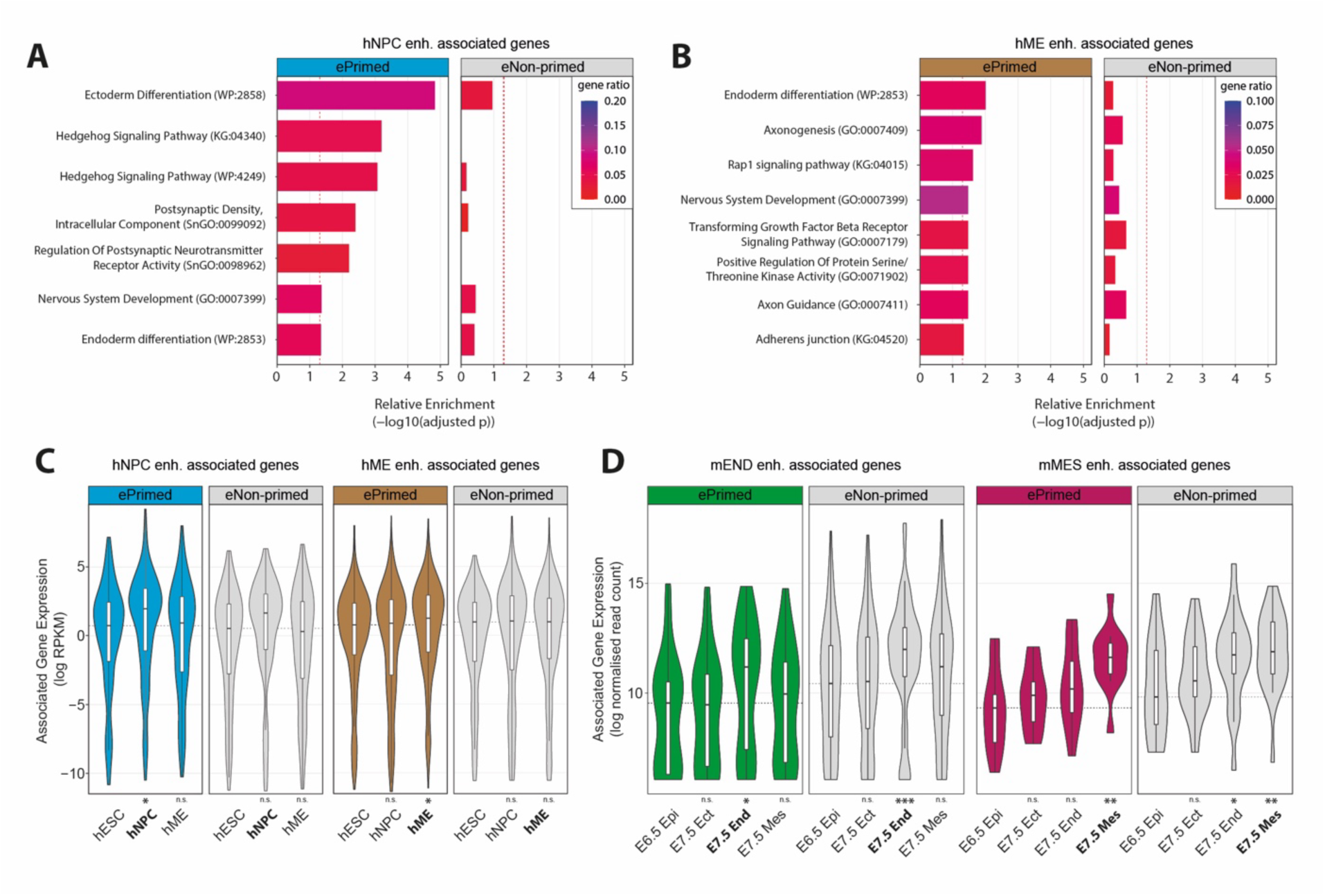
**(A-B)** Gene ontology enrichment analysis for ePrimed and eNon-primed enhancer-associated genes as defined by PCHiC/proximity hybrid model. Significance threshold of p<0.05 is displayed (red dashed line). **(A)** Analysis of ePrimed and eNon-primed hNPC enhancer-associated genes and **(B)** ePrimed and eNon-primed hME enhancer-associated genes. **(C-D)** Overlaid box and violin plots show expression of enhancer-associated genes. Box plots show median levels and the first and third quartile, whiskers show 1.5× the interquartile range. Results from Welch two-sample t-test against E6.5 epiblast/hESC are displayed (p<0.01 **, p<0.05 *). **(C)** Expression (log RPKM) within *in vitro* RNA-seq data collected from hESC, hNPC, hME cultures of genes associated exclusively with either ePrimed hNPC (blue) enhancers, eNon-primed hNPC (grey) enhancers, ePrimed hME (brown) enhancers, or eNon-primed hME (grey) enhancers. Median expression for each gene group within hESCs is also shown (grey dashed line). **(D)** Expression (log normalized counts) within *in vivo* scNMT-seq data collected from E6.5 epiblast and E7.5 ectoderm/ endoderm/ mesoderm tissues for genes associated exclusively with either ePrimed E7.5 mEND (green) enhancers, eNon-primed E7.5 mEND (grey) enhancers, ePrimed E7.5 mMES (purple) enhancers, or eNon-primed E7.5 mMES (grey) enhancers. Median expression for each gene group within E6.5 epiblast is also shown (grey dashed line).

Next, we analysed the expression dynamics of these gene groups during human *in vitro* differentiation and mouse *in vivo* gastrulation. In the hESC differentiation model, genes exclusively associated with ePrimed hNPC enhancers were significantly upregulated (1.9 fold) upon differentiation into hNPCs (Figure 2C). This upregulation appears to be lineage-specific to hNPC wherein the corresponding hNPC enhancer becomes marked by the active mark H3K27ac (Figure 1A), with no observed upregulation upon differentiation into hMEs where the corresponding enhancer is not activated (Figure 1A, 2C). Likewise, genes associated with primed hME enhancers are upregulated specifically in hME lineage differentiation (Figure 2C). Comparable lineage-specific expression patterns were observed for genes associated with ePrimed enhancers during mouse gastrulation (Figure 2D). Genes associated with ePrimed E7.5 mEND enhancers are upregulated specifically in E7.5 endoderm (Figure 2D) coinciding with when the corresponding enhancer gains H3K27ac (Figure 1B). This is specific to endoderm as the same patterns are not seen within E7.5 mesoderm or ectoderm tissues where the corresponding ePrimed E7.5 mEND enhancers do not gain these active marks (Figure 1B, 2D). Similarly, genes associated with ePrimed mMES enhancers are significantly upregulated exclusively within the mesoderm at E7.5 (Figure 2D). Conversely, the regulation of eNon-primed enhancer-associated gene groups is far less consistent, with many showing less lineage-specificity in their expression. In the hESC differentiation model, genes exclusively associated with eNon-primed hNPC enhancers show an observable but statistically insignificant upregulation within hNPCs (Figure 2C). In contrast, genes associated with eNon-primed hME enhancers show no expression changes upon differentiation from hESCs into hNPCs or hMEs (Figure 2C). Some eNon-primed enhancer-associated genes do show lineage-specific expression, with eNon-primed E7.5 mEND enhancer-associated genes displaying significant upregulation exclusively with mEND lineage at E7.5 (Figure 2D). However, genes associated with eNon-primed E7.5 mMES enhancers are not lineage-specific in their expression and are significantly upregulated within both mesoderm and endoderm E7.5 tissues (Figure 2D). Notably, we failed to detect significant upregulation of E7.5 mECT enhancer-associated genes, possibly as E6.5 epiblast already exists within a partial ectodermal state (Supp. Figure 6C). Interestingly, within the majority of somatic tissues eNon-primed enhancer-associated genes were typically upregulated relative to ePrimed enhancer-associated genes, suggesting that they are enriched for non lineage-specific post-gastrulation processes (Supp. Figure 7). These observations suggest that the epigenetic priming signature of enhancers within the epiblast is associated with an increased lineage-specificity in the regulation of associated gene networks, consistent with a central role for enhancer priming in the regulation of key developmental gene networks.

### A subset of early brain lineage-specific enhancers are epigenetically primed in the epiblast

Curiously, we found that only a small proportion (<1% in mouse & human) of ePrimed enhancers become active enhancers (i.e. gain H3K27ac) within tissues immediately derived from the epiblast. To understand if enhancer priming was also found at enhancers associated with later cell fate decisions, we next analysed lineage-specific enhancers only activated at a significantly later embryonic developmental timepoint. Specifically, we focused on comparative neural tissues collected at E11.5 from mouse embryos and 7 weeks post conception (7pcw) tissues from human embryos. Peak calling of H3K27ac ChIP-seq data collected from E11.5 mouse forebrain identified 5,583 distal peaks. These putative active enhancer elements were further filtered for lineage-specific enhancers by removal of sites with H3K27ac signal in earlier developmental tissues, which yielded 839 E11.5 forebrain-specific enhancers. As validation of our enhancer calling, we compared these enhancer sites to those within the VISTA Enhancer Browser (21). Out of 183 mouse enhancers annotated with forebrain activity within the VISTA Enhancer Browser dataset, 155 overlap our putative E11.5 forebrain enhancers, 8 of which overlap enhancers that are active exclusively within the forebrain (Supp. Fig 8A-B). Similarly, we called 77,235 distal H3K27ac peaks within human 7pcw fetal brain tissues, 51,187 of which were lineage-specific, with no evidence of H3K27ac signal within the aforementioned earlier tissues analysed. Strikingly, there are 4,218 7pcw fetal brain ePrimed enhancers (Figure 3A,C) and 211 E11.5 forebrain ePrimed enhancers (Figure 3B,D) that are found to be in an epigenetically primed state (i.e. H3K4me1 positive) already within the epiblast. These later-development ePrimed enhancers show similar H3K4me1 signals in the absence of poised enhancer-associated H3K27me3 or active enhancer-associated levels of H3K27ac. Similar to what we observed at the other ePrimed enhancers described above there is a weak H3K27ac signal at a subset of 7pcw fetal brain and E11.5 forebrain ePrimed enhancers, although the signal is much lower than at active hESC/mESC enhancers (Figure 3C,D). Furthermore, E11.5 forebrain and 7pcw fetal brain ePrimed enhancers were also associated with decreased DNA methylation and increased chromatin accessibility (Supp. Figure 9A-D). Gene ontology analysis of E11.5 forebrain and 7pcw fetal brain ePrimed enhancer-associated genes showed a significant enrichment for key nervous system developmental processes and neurogenesis (Figure 3E,F). In contrast, eNon-primed enhancer-associated genes were generally not enriched for neurodevelopmental processes (Figure 3E,F). Overall, our findings suggest that epigenetic priming not only occurs at enhancers that will be activated in immediately following developmental stages, but also at enhancers that become active at much later stages.

**Figure 3.**
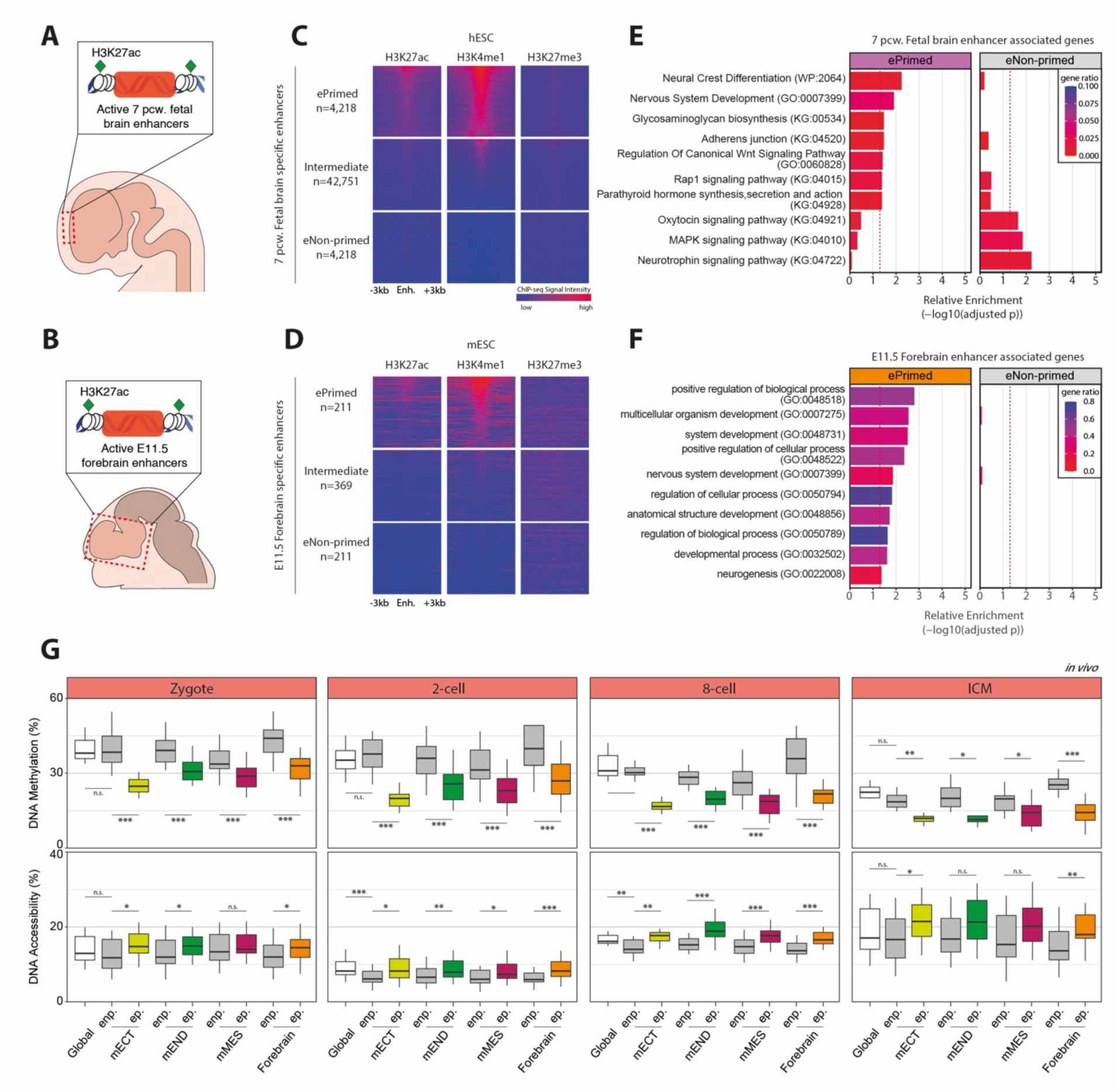
**(A-B)** Graphical representations of tissue dissections taken for H3K27ac ChIP-seq libraries for the identification of active enhancers within **(A)** human 7 post conception week fetal brain and **(B)** mouse E11.5 forebrain. **(C)** Heatmaps of ChIP-seq H3K27ac, H3K4me1 and H3K27me3 data within hESC cells in culture at lineage-specific enhancers for 7pcw fetal brain tissues that are called as ePrimed, eNon-primed and intermediate states within the hESCs. **(D)** Heatmaps of ChIP-seq H3K27ac, H3K4me1 and H3K27me3 data within mESC cells in culture at lineage-specific enhancers for E11.5 forebrain tissues that are called as ePrimed, eNon-primed and intermediate states within the mESCs. **(E-F)** Gene ontology enrichment analysis for **(E)** ePrimed and eNon-primed 7pcw fetal brain enhancer-associated genes, and **(F)** ePrimed and eNon-primed E11.5 forebrain enhancer-associated genes. Significance threshold of p<0.05 is displayed (red dashed line). **(G)** COOL-seq analysis of DNA methylation and chromatin accessibility levels within preimplantation mouse embryos over 500bp core of E7.5 mECT, E7.5 mEND, E7.5 mMES and E11.5 forebrain called enhancer sub-groups: ePrimed (ep.) and eNon-primed (enp.). Global levels for each cell is shown (white). Results for comparison of means between groups (ANOVA) is shown (p<0.001 ***, p<0.01 **, p<0.05 *).

### Epigenetic priming of enhancers originates in the zygote

To study when epigenetic priming of ePrimed enhancers is first established, we analysed single-cell multi-omics sequencing data from initial stages of mouse embryogenesis ranging from the zygote to the early blastocyst captured by COOL-seq, a sequencing technology that can simultaneously analyse chromatin accessibility, DNA methylation, and DNA copy number variation for individual mammalian cells (22). Surprisingly, we observed that E7.5 germ-layer and E11.5 forebrain ePrimed enhancers were hypomethylated as early as the fertilised zygote, and remained hypomethylated in the inner cell mass of the blastocyst (Figure 3G). Similarly, we found that chromatin at ePrimed enhancers is more accessible than at eNon-primed enhancers in the zygote, which further increases upon establishment of the ICM (Figure 3G).

As reduced levels of DNA methylation marks were consistently observed at primed enhancer sites, we next looked at the dependence of this pattern on the enzymes that are involved in the establishment and removal of DNA methylation marks. DNA methyltransferases (DNMTs) are responsible for *de novo* methylation (DNMT3a/DNMT3b) (23), while ten-eleven translocation proteins (TETs) facilitate the removal of DNA methylation marks by oxidation (24–26). We analysed DNA methylation levels of hESC cell lines with combinations of genetic knockouts for DNMTs and TETs (27). In DNMT3A/DNMT3B double knockouts, primed enhancers show a significant decrease in DNA methylation (Supp. Figure 9E). In TET1/2/3 triple knockout cells, there is a loss of the hypomethylation signature at primed enhancers, which is then recovered when DNMT3A/DNMT3B are knocked out simultaneously (Supp. Figure 9E). These results suggest that the DNA hypomethylation at ePrimed enhancers in hESCs is maintained by TET/DNMT antagonistic activity, similar to the dynamics and antagonistic activity that occurs at active enhancers but not at eNon-primed enhancer sites (Supp. Figure 9E).

Argelaguet et al. (16), showed that primed ectoderm lineage enhancers exhibit DNA hypomethylation in non-ectodermal germ layers, suggesting an epigenetic memory of the priming event. We observed a similar epigenetic memory of the priming event within our ePrimed enhancer groups, with DNA hypomethylation persisting in germ layer tissues in which the enhancer had not been activated. For example, ePrimed mEND enhancers remained hypomethylated when cells differentiated into ectoderm and mesoderm (Supp. Figure 9F). Notably, we observed that the hypomethylation signature at ePrimed enhancers persisted into adult somatic tissues, similarly regardless of the germ-layer origin of the tissue (Supp. Figure 10).

### Defining sequence determinants of ePrimed enhancers using naturally occurring regulatory human genetic variation

Having identified groups of ePrimed and eNon-primed enhancers, we next asked if the DNA sequence composition of primed enhancers could contribute to their epigenetic priming. Since previous studies have shown associations with H3K4me1 marks and the binding of pioneer factors (28–30) we performed motif enrichment analysis. This revealed an enrichment of various TF motifs within ePrimed 7pcw fetal brain enhancer regions when compared to eNon-primed counterparts. The enriched motifs include expected neural development-associated TFs, such as MAZ (31) and SOX6 (31) (Figure 4A). The ePrimed enhancer enriched motif sequences also include motifs for TFs which are more highly expressed at epiblast-like stages, such as SOX3, SOX2 and ZIC3, which could be involved in the establishment or maintenance of the primed state (Figure 4A). However these motifs are not exclusive to ePrimed enhancers as they are also found prevalent within their respective non-primed counterparts (Figure 4A).

**Figure 4.**
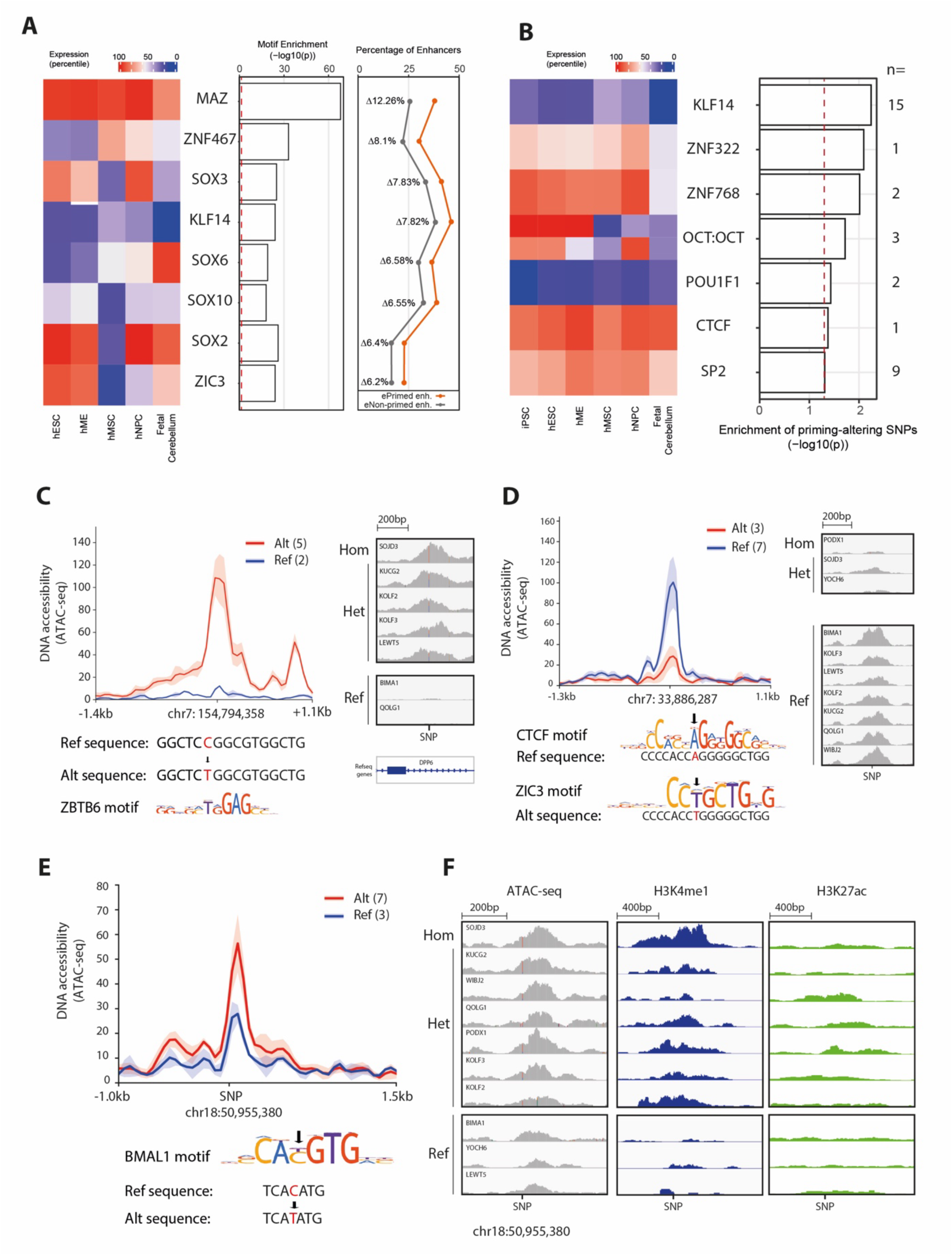
**(A)** Motif enrichment analysis of human ePrimed vs eNon-primed 7pcw fetal brain enhancers. Heatmap of relative expression (percentile rank) of the significantly enriched motif elements within hESC, hME, hMSC, hNPC cultures and *in vivo* fetal cerebellum tissue (collected 72-129 post conception days). Barplot shows p-values for motif enrichment test with significance threshold of p<0.05 displayed (red dashed line). Line plot shows percentage of enhancers which contain an instance of each motif for ePrimed (orange) and eNon-primed (grey) 7pcw fetal brain enhancer sub-groups. Points are annotated with the percentage differential between ePrimed and eNon-primed sub-groups. **(B)** Motifs associated with priming-altering SNPs within HipSci iPSC lines. Heatmap of relative expression (percentile rank) of the significantly enriched motif elements within hESC, hME, hMSC, hNPC cultures and *in vivo* fetal cerebellum tissue. Expression of OCT4 and OCT6 is shown for OCT:OCT. Barplot shows p-values of association test between disruption of priming and presence of SNP within the motif sequence (Fisher’s exact test) with significance threshold of p<0.05 displayed (red dashed line). **(C-E)** Profiles of chromatin accessibility and genome browser visualisation of priming-altering associated SNP examples within ePrimed 7pcw fetal brain enhancers. HipSci donors are grouped based on the sequence variant. Running averages in 50bp windows around the SNP are shown. Solid lines show the mean across donor lines and shaded areas represent the standard deviation. Genome browser visualization of ATAC-seq (grey) signal within the donor lines, grouped by allelic presence of SNP. **(C)** Example of introduction of ZBTB6 motif by C→T SNP at chr7:154,794,358 (hg38). Donor lines containing a second confounding SNP were excluded. **(D)** Example of altering CTCF motif to ZIC3 motif by A→T SNP at chr7:33,886,287 (hg38). **(E)** Example of altering a BMAL1 motif by C→T SNP at chr18:50,955,380. **(F)** Genome browser visualization of ATAC-seq (grey), H3K4me1 (blue) and H3K27ac (green) signal within the donor lines, grouped by allelic presence of SNP chr18:50,955,380 (hg38).

We next used human induced pluripotent stem cell lines from different donors to functionally interrogate TF motif sequences and their respective importance for the observed epigenetic priming signatures. The human induced pluripotency stem cell initiative (HipSci) has derived induced pluripotent stem cells (iPSC) from skin fibroblasts taken from a number of patients and healthy donors containing a collection of germline inherited and somatically acquired polymorphisms (32,33). This naturally occurring genetic variation can be used to query the relationship between primary sequence and enhancer priming. We analysed whole genome sequencing datasets and performed ATAC-seq (34) and CUT&Tag (35) for H3K4me1 and H3K27ac in 10 HipSci iPSC lines, representing 9 unique healthy donors. Many of our primed 7pcw brain enhancers coincided with chromatin accessibility peaks and H3K4me1 signal in the absence of H3K27ac, suggesting that these iPSC lines recapitulate the priming state observed in the epiblast at many of these enhancers (Supp. Figure 11A-B). We next identified single nucleotide polymorphisms (SNPs) which overlapped TF motifs in these primed enhancers and identified SNPs that disrupted their priming by identifying either a gain or loss of ATAC-seq peak when a SNP was present. For 145 different TF motifs overlapping a variant in an enhancer with altered accessibility, 7 TF motifs were significantly enriched (KLF14, ZNF322, ZNF768, OCT:OCT, POU1F1, CTCF, SP2) (Figure 4B).

We observed three different scenarios of how particular SNPs can alter TF motif sequences. First, we found instances where SNPs would introduce/ablate a motif entirely, such as the introduction of a ZBTB6 motif which coincided with a marked increase in priming of the associated enhancer (Figure 4C). Second, we observed instances where a SNP would alter a key residue within a motif and result in the introduction of an alternative TF motif, such as the replacement of a CTCF motif for a ZIC3 motif which resulted in a loss of associated priming state (Figure 4D). Lastly, we observed instances where a SNP appears to alter the preferred base within the motif towards a less preferred base, potentially weakening the binding affinity. One example was the change of a preferred C residue to a weaker affinity T base within the motif for the BMAL1 pioneer-like TF factor; this potential weaker affinity motif was associated with an increase of priming signature at the associated enhancer, with both increased DNA accessibility and H3K4me1 signal (Figure 4E-F). This could be caused by preferences in degenerate motifs for pioneer factors when targeting nucleosome bound DNA (31). Our results demonstrate that while motif enrichment analysis is unable to identify individual sequences that are predictive for enhancer priming, naturally occurring genetic variation can be integrated with epigenetic profiling to reveal potentially important enhancer motifs.

## Discussion

This study has identified a program of enhancer priming in human embryo development similar to that previously observed associated with mouse gastrulation (16) indicative of an evolutionarily conserved mammalian program for epigenetic priming of lineage-specific regulatory elements. Priming signatures are more pronounced in ectoderm lineage enhancers, especially in late epiblast tissues, however our findings suggest enhancer priming is used to also establish the capacity for differentiation into the mesoderm and endoderm fates. Similar to enrichments of poised enhancer-associated gene networks (8), this conserved program of enhancer priming appears to be linked with the lineage-specific regulation of key developmental gene networks, pointing to an important role in healthy embryonic development.

What remains unclear is the functional impact of epigenetic priming of these enhancers, whether their priming is required for accurate and rapid activation of associated genes to control developmental gene networks as well as the functional importance of the priming associated epigenetic marks themselves. While the H3K4me1 mark identifies these pre-active state enhancers, recent studies have begun to question if the mark itself or associated histone modifiers are responsible for driving enhancer activation and subsequent transcriptional responses. Many enhancer elements, including key regulators of early differentiation, still gain H3K27ac upon knockout of H3K4me1 histone modifiers MLL3 and MLL4 (36), however conversely many enhancers lose the ability to become activated upon MLL4 KO (37). These conflicting results suggest a potential context dependency for the importance of H3K4me1 and its writers. However these KO studies are unable to distinguish between the importance of the H3K4me1 mark and potential enzymatic activity-independent functions of its writers who themselves may recruit the transcriptional machinery directly.

Questions also remain regarding the temporal order and interplay between the various epigenetic marks associated with the primed enhancer state. Interestingly, our observations reveal that primed enhancers are hypomethylated and show nucleosome displacement as early as the zygote. It is therefore possible that these marks precede, and may be required to recruit the observed H3K4me1 signature. DNA hypomethylation is indeed observed developmentally early, in the zygote, and may facilitate the recruitment of TFs that subsequently recruit H3K4 methyltransferases (38–41). Conversely it has been hypothesised that the H3K4me1 reader, TIP60, can promote the integration of H2A.Z containing nucleosomes to create more dynamic and open chromatin structures (42,43). These hypotheses could also point towards a positive feedback loop between these epigenetic layers, and may explain why the priming signature is stable enough to persist for long periods of development and many cell divisions as observed in our models and previous observations of stable H3K4me1 marked regions during zebrafish development (44). Capture and analysis of H3K4me1 datasets of earlier developmental time points would be required to dissect the relative timings of these epigenetic marks, while targeted epigenetic manipulation of ePrimed and eNon-primed enhancers in differentiation models in combination with inducible degron *in vivo* models could help dissect the respective roles and interplay between these epigenetic marks.

Another important question is how priming of the particular enhancer regions is initiated during development. Our study identified a number of enriched sequence motifs but no universal priming-associated enhancer sequence signature. A limitation of our study is that the analysis has not considered the relative strength of the motifs present or whether combinations of motifs may predict sites which undergo epigenetic priming. Furthermore the identified enriched motifs will require further experimental interrogation to validate their functional role. Additionally this analysis is limited by the natural variation within the donor lines used; the power of this analysis could be further increased by the inclusion of additional lines and therefore increased SNP instances within the enhancer regions of interest. The inability to identify singular priming-associated motif sequences could suggest that enhancer priming is a complex process, defined by various sequence signatures in concert interplaying with additional factors such as the enhancer’s location within the genome. In this context, it is worth noting that enhancers have been shown to tolerate substantial sequence changes (45).

Interestingly, hypomethylation “scars” persist at primed enhancers within adult somatic tissues. Previous studies have shown that developmental enhancers with hypomethylation scars retain the capacity to be reactivated upon PRC2 disruption (46), and suggested that reactivation of developmental enhancers facilitates the regeneration of tissues in response to damage (47). It is possible that hypomethylation scars at primed enhancers provide an epigenetic memory of developmental cell states, facilitating the return to these states upon perturbations required for the repair or rejuvenation of tissues. This would provide an interesting model to interrogate the importance of enhancer priming and maintenance of their epigenetic memory for the capacity of somatic cells to rejuvenate in perturbations such as Yamanaka factor induced cellular rejuvenation (48).

## Conclusion

In this study, we perform multi-omic profiling of human and mouse embryonic development to determine the dynamics of enhancer epigenetic priming and to investigate its role in regulating cell fate decisions. We further leverage naturally occurring genetic variation in human pluripotent stem cell lines to identify sequence determinants and potential trans-acting factors that establish the primed state at key human developmental enhancers. Our work provides evidence of a conserved program of epigenetic priming for lineage-specific enhancers that exists in early human embryo development, and demonstrates that epigenetic priming is present at enhancer elements for all three germ layers in both human and mouse epiblast tissues. The results of this study suggest that the observed epigenetic priming of enhancers is associated with the lineage-specific regulation of key developmental gene networks. Our findings that epigenetic priming within the epiblast also occurs at enhancer elements specific for later developmental stages (E11.5 in mouse, and 7 weeks post conception in human) also suggest that enhancer priming occurs far earlier than previously anticipated, and that its functional consequences manifest through remarkably long developmental periods along cell specification trajectories.

## Methods

### Human iPSC Cell culture

Human induced pluripotent stem cell (hiPSC) lines (HPSI1113i-bima_1, HPSI0114i-kolf_2, HPSI0114i-kolf_3, HPSI0214i-kucg_2, HPSI0514i-letw_5, HPSI1113i-podx_1, HPSI1113i-qolg_1, HPSI0314i-sojd_3, HPSI0214i-wibj_2, and HPSI0215i-yoch_6) were purchased from the Human Induced Pluripotent Stem Cells Initiative (HipSci; https://www.hipsci.org/). All cell lines were cultured in TeSR-E8 media complete with supplement (Stemcell Technologies; 05990) on plates coated with Vitronectin (Thermo Fisher Scientific; A14700) at 37°C under 5% CO_2_ and normal O_2_ levels. Complete media was refreshed every 24 hours. For the first day after revival, 10 µM Rho-associated protein Kinase (ROCK) inhibitor (Cell Guidance Systems; Y-27632) was added to the media. Cells were passaged at ratios ranging from 1/6 to 1/8 using 0.5 mM EDTA (Life Technologies; AM9260G) upon reaching approximately 70% confluency. Regular tests ensured that all HiPSCs lines were negative for mycoplasma contamination.

### ATAC-seq

ATAC-seq libraries were generated as described in Corces et al.(49), with minor modifications. After washing once with 1X DPBS (Life Technologies; 14190144), hiPSCs were harvested with accutase (StemCell Technologies; 07922) by incubation for 5 min at 37°C. After accutase neutralisation with TeSR-E8 medium cells were centrifuged at 300 x g for 3 min at RT. After resuspension in 1X DPBS, 50,000 cells underwent centrifugation at 500 x g for 5 min at 4°C in a swing arm rotor centrifuge, then lysis in 50µl of cold ATAC Resuspension Buffer (10mM Tris-HCl pH 7.5, 10mM NaCl, 3mM MgCl_2_) containing 0.1% IGEPAL-630, 0.1% Tween-20, and 0.01% digitonin for 3 min on ice. The lysate was then topped up with 1ml cold ATAC Resuspension Buffer containing 0.1% Tween-20 and centrifuged at 500 x g for 10 min at 4°C in a swing arm rotor centrifuge. The pellet was then resuspended in 50µl of transposition mixture (1X Illumina Tagment DNA Buffer and 100nM Illumina TDE1 Tagment DNA Enzyme transposase (Illumina; 20034197), 0.33X DPBS, 0.1% Tween-20, 0.01% digitonin) and incubated at 37°C for 30 min with 1000 RPM mixing.

After undergoing purification with the Zymo DNA Clean & Concentrator-5 kit (Zymo Research; D4003), samples were eluted in 21µl of elution buffer, and combined with 2.5µl of 25µM i5 primer, 2.5µl of 25µM i7 primer, and 25µl 2X NEBNext High-Fidelity 2X PCR Master Mix (NEB; M0541S). A PCR was then performed with the following conditions: 72°C for 5 min, 98°C for 30 sec, 8 cycles of [98°C for 10 sec, 63°C for 30 sec, 72°C for 1 min]. The samples then underwent a second Zymo DNA Clean & Concentrator-5 purification and were eluted in 30µl of elution buffer prior to a 1.2X AMPure XP bead (Beckman Coulter; A63881) cleanup. Following QC on a Bioanalyzer, libraries were multiplexed and sequenced (paired-end 50bp) using a HiSeq 2000 instrument (Illumina).

### CUT&Tag

CUT&Tag libraries were generated according to the EpiCypher protocol (35) with minor modifications. As described for ATAC-seq, hiPSCs were harvested with accutase (StemCell Technologies; 07922). After resuspension in 1X DPBS (Life Technologies; 14190144), 2 x 10^5^ cells per CUT&Tag reaction underwent centrifugation at 300 x g for 3 min at RT. Cells were then lysed in 100μl of cold Nuclear Extraction (NE) Buffer (20mM HEPES–KOH, pH 7.9, 10mM KCl, 0.1% Triton X-100, 20% Glycerol, 0.5mM Spermidine (Sigma-Aldrich; 05292), 1x cOmplete^TM^, Mini, EDTA-free Protease Inhibitor (Roche; 11836170001)) per CUT&Tag reaction for 10 min on ice. The nuclei were then centrifuged for 3 min at 600 x g at RT before resuspension in cold NE Buffer to achieve a final concentration of 1.2×10^6^ nuclei/ml.

For each CUT&Tag reaction, 11μl of concanavalin A (ConA) beads (EpiCypher; 21-1401) were washed twice on a magnetic stand with 100 μl of Bead Activation (BA) Buffer (20mM HEPES, pH 7.9, 10mM KCl, 1mM CaCl_2_, 1mM MnCl_2_). The beads were then resuspended in 11μl of BA Buffer, and 10μl beads were aliquoted into a 0.2ml tube. Each tube of activated ConA beads was then incubated with 100μl of nuclei for 10 min at RT. The supernatant was then removed with a magnet and the beads were resuspended in 50μl of cold Antibody150 Buffer (20mM HEPES, pH 7.5, 150mM NaCl, 0.5mM Spermidine, 1x cOmplete^TM^, Mini, EDTA-free Protease Inhibitor, 0.01% digitonin (Sigma-Aldrich, D141-100MG), 2mM EDTA) containing a 1:50 dilution of rabbit primary antibody (anti-H3K4me1 (Active Motif; 39298) or anti-H3K27ac (Abcam; ab4729)) and incubated overnight at 4°C on a rocker.

The following day, the supernatant was discarded with a magnet and the beads were incubated in 50μl of Digitonin150 Buffer (20mM HEPES, pH 7.5, 150mM NaCl, 0.5mM Spermidine, 1x cOmplete^TM^, Mini, EDTA-free Protease Inhibitor, 0.01% digitonin) containing 0.5μg of anti-rabbit secondary antibody (Epicypher; 13-0047) for 1 hour at RT. The beads were then washed twice on a magnet with Digitonin150 Buffer, then incubated in 50μl of Digitonin300 Buffer (20mM HEPES, pH 7.5, 300mM NaCl, 0.5mM Spermidine, 1x cOmplete^TM^, Mini, EDTA-free Protease Inhibitor, 0.01% digitonin), and 1.25μl of CUTANA pAG-TN5 (EpiCypher; 15-1017) for 1 hour at RT. The beads were then washed twice with a magnet by resuspension in Digitonin300 Buffer, then incubated in 50μl of chilled Tagmentation Buffer (Digitonin 300 Buffer, 10mM MgCl_2_) for 1 hour at 37°C. The supernatant was then discarded with a magnet and the beads were washed once with 50μl RT TAPS Buffer (10mM TAPS, pH 8.5, 0.2mM EDTA). The supernatant was again removed with the magnet and the beads were resuspended in 5μl of RT SDS Release Buffer (10mM TAPS, pH 8.5, 0.1% SDS) and vortexed on maximum speed for 10 sec followed by a brief centrifugation. The beads were then incubated for 1 hour at 58°C before the addition of 15μl of RT SDS Quench Buffer (0.67% Triton-X 100 in Molecular grade H_2_O), vortexing at maximum speed for 10 sec, and a brief centrifugation.

For library amplification, samples were combined with 2μl of 5µM i5 primer, 2μl of 5µM i7 primer and 25μl of CUTANA High Fidelity PCR mix (EpiCypher; 15-1018). The following PCR programme was then used for all libraries: 58°C for 5 min, 72°C for 5 min, and 98°C for 45 sec. This was followed by 11 cycles for anti-H3K27ac and 13 cycles for anti-H3K4me1 of [ 98°C for 15 sec and 60°C for 10 sec], before a final extension at 72°C for 1 min. Libraries then underwent two 1X AMPure XP bead (Beckman Coulter; A63881) cleanups. Following QC on a Bioanalyzer, libraries were multiplexed and sequenced (paired-end 150 bp) using a NovaSeq 6000 instrument (Illumina).

### Human preimplantation embryo scNMT-seq

The use of human embryos for this work has been approved by the Multi-Centre Research Ethics Committee and licensed by the Human Fertilization and Embryology Authority of the United Kingdom, under Research License R0178. Embryos from embryonic days 5-6 were collected in Corujo-Simon et al. (50) in which scRNA-seq was performed using the RNA half of the scNMT-seq method (51). In this study we used the gDNA from the same cells to generate DNA methylation and chromatin accessibility data using the published scNMT-seq protocol (52). Briefly, genomic DNA was purified using AMPure XP beads then bisulfite converted with the Zymo EZ-96 DNA Methylation-Direct MagPrep kit. Sequencing libraries were then prepared from bisulfite converted fragments via random primed synthesis using oligos containing Illumina primer sequences, followed by indexing PCR. Pooled libraries were sequenced using an Illumina HiSeq 2000 instrument using 125bp paired-end reads (day 5 and day 6 embryos). Fastq files were aligned to the GRCh38 build of the human genome and CpG methylation and GpC accessibility files generated as previously described (16).

### Defining Lineage-specific enhancer states within the epiblast

Murine ChIP-seq data for mESC *in vitro* cultures were obtained from Parry et al. 2023 (GSE223565), *in vivo* E7.5 tissues from GSE125318, and *in vivo* fetal tissues from the ENCODE project (53). Reads were trimmed using Trim Galore v0.6.1 (using Cutadapt v1.18) (54) and mapped to *Mus musculus* GRCm38 using Bowtie2 v2.4.1 (55). Human ChIP-seq data for Primed hESC *in vitro* cultures were obtained from SRP000941, and for *in vivo* 7pcw fetal brain tissue from GSE63648. Reads were trimmed using Trim Galore v0.6.1 (using Cutadapt v1.18) and mapped to *Homo sapiens* GRCh38 using Bowtie2 v2.4.1 (55). Quantification of these marks was performed using SeqMonk v1.48.2 (56). H3K27ac peaks were called using integrated MACS pipeline (p-value cutoff 1.0E-5).

For early development human enhancers (hME,hMSC,hNPC), lineage-specific enhancers were obtained by excluding previously annotated enhancers from Xie et al.(17), and intersected with called hESC, hME, hMSC, and hNPC H3K27ac peaks using BEDtools intersect (57), subsetting for enhancer instances that failed to overlap H3K27ac peaks in tissues outside of their specific lineage. 7pcw fetal brain lineage-specific enhancers were defined as 7pcw fetal brain H3K27ac peaks that do not overlap with an annotated TSS (±200bp) or an ulterior-lineage H3K27ac peak (hESC,hMSC,hME,hNPC). For mouse enhancers, E7.5 mECT/mEND/mMES lineage-specific enhancers were defined as respective H3K27ac peaks that do not overlap with an annotated TSS (±200bp) or an ulterior-lineage H3K27ac peak. E11.5 lineage-specific enhancers were similarly defined as respective H3K27ac peaks that do not overlap with an annotated TSS (±200bp) or an ulterior E11.5, E7.5 or mESC lineage H3K27ac peak. logRPKM counts for ChIP-seq reads were quantified over 1500bp probe centred on the enhancer region. Enhancer states within human epiblast were sequentially defined using these counts: Active enhancers as regions with H3K27ac >2.97, Poised as remaining regions with H3K27me3 >2, Primed as remaining regions with H3K4me1 >1.2, epiblast-Inactive as all remaining regions. Enhancer states within mouse epiblast were similarly defined as active enhancers with H3K27ac >1.2, poised with H3K27me3 >1.25, primed with H3K4me1 >-0.4 , and epiblast-inactive for remaining regions. For visualisation of ChIP-seq, signal heatmaps were generated using Homer tools v4.11 (28) and Samtools v1.11 (58).

### Chromatin Accessibility and DNA methylation at enhancers

ATAC-seq data for mESC *in vitro* cultures were obtained from GSE81679, whole genome bisulfite (WGBS) data from DRA003471. Reads were trimmed using Trim Galore v0.6.6 (using Cutadapt v2.3) and mapped to *Mus musculus* GRCm38 using Bismark v0.23.0 (59). ATAC-seq data for hESC cultures were obtained from GSE101074, and WGBS data from GSE75868. WGBS data for hESC cultures with TET and DNMT combinatorial knockouts were obtained from GSE126958. WGBS data for human somatic tissues were obtained from the NIH Roadmap Epigenetics Consortium (60). Reads were trimmed using Trim Galore v0.6.6 (using Cutadapt v2.3) and mapped to *Homo sapiens* GRCh38 using Bismark v0.23.0. For visualisation of ATAC-seq signals, Homertools v4.11 annotatepeaks function was used over a ±3kb region centred on the enhancer region. Quantitation of DNA methylation *in vitro* datasets was performed using Seqmonk Bisulfite methylation quantitation function over a 500bp probe centred on the enhancer. Global levels of methylation calculated as the mean of 10kb running windows. Analysis of scNMT and COOL-seq data was performed using methods and scripts outlined in Argelaguet et al. (16).

### Analysis of enhancer-associated gene networks

Significant Promoter Capture Hi-C interactions within mESCs were obtained from GSE223578, and within hESCs from GSE86821. Enhancers overlapping promoter-interacting fragments were identified by BEDtools intersect; proximal promoters within 20kb of enhancers were also considered potentially interacting. Promoters were filtered for those interacting with enhancers from a singular lineage and enhancer-state. Gene ontology enrichments were performed using the enrichr R package v3.2 (github.com/wjawaid/enrichR). Gene expression data for hESC, hME, hMSC, hNPC *in vitro* cultures were obtained from SRP000941, for human fetal tissues from GSE156793, and for human somatic tissues from GSE144530. Expression of protein-coding genes was converted to percentile position within the tissue to facilitate visualisation across tissues and gene expression data types.

### Identification of priming-altering SNPs within human iPSCs

ATAC-seq peaks were called separately for each replicate by MACS2 (61) (options: -q 0.01 -- nomodel --keep-dup all ); high confidence peaks were called as peaks present within at least two of the three replicates. SNPs and indels within the 10 HipSci donors were called using GATK (62). BEDtools intersect was used to identify 7pcw fetal brain enhancers which overlapped SNPs or ATAC peaks in any donor. High-confidence accessible enhancers were called as overlapping ATAC peaks in at least two of the three replicates for each donor line. Known motifs were called in these enhancers using Homer v4.11. Enrichment for TF motifs within enhancers coinciding with altered accessibility when a SNP is present was determined using a Fisher’s exact test. For visualisation of ATAC signal at interesting sites, deeptools v3.5.1 was used (63). Donors were grouped into their genotype on a per SNP basis, then counts within the enhancer of interest were extracted and compressed to generate one bigwig per genotype using bedGraphToBigWig (63,64). These bigwigs were input into deeptools computeMatrix and plotProfile with mean and standard deviation plotted.

For visualisation of Cut&Tag signal across HipSci donor lines, initial quality check of paired-end reads was performed by FastQC v0.11.5 (65). Nextera Transposase adapter and low-quality bases were eliminated using Cutadapt v1.17 (66). Reads were mapped to the hg38 reference using bowtie2 v2.3.4.2, mapped pairs with mapping quality <30 were removed. Replicates were removed using GATK Picard MarkDuplicates. Genome browser tracks in bigwig format were generated using deepTools bamCoverage (options: binsize = 10, RPGC normalization).

## Declarations

### Ethics Approval and consent to participate

Use of human tissues approved by the Multi-Centre Research Ethics Committee and licensed by the Human Fertilization and Embryology Authority of the United Kingdom, under Research License R0178.

### Consent for publication

Not applicable

### Availability of data and materials

Datasets will be made available on GEO upon publication.

### Authors’ contributions

CDT, SS & WR designed and conceived the study. JP performed pilot analysis. CDT & JI performed most of the bioinformatics analysis. ATAC-seq experiments were performed by SB. CUT&Tag experiments were performed by FT & OC. Primary analysis of HipSci CUT&Tag and RNA-seq datasets was performed by UG & OD. Human embryonic tissues were prepared by JN, TL & FVM. SC performed scNMT-seq, and analysis performed by SC & RAC. IK & AS provided help and suggestions in data interpretation. The paper was written by CDT with supervision from WR & SS, along with contributions and suggestions from all other authors.

## Acknowledgements

We thank Peter Rugg-Gunn for expert advice and help with human iPSC culture, and Dominik Muehlen for expert advice on CUT&Tag. We also wish to thank the Babraham Institute Bioinformatics group for their support and all members of the Reik lab and the Babraham Institute Epigenetics Programme for their scientific discussion.

---

**Supp. Figure 1.**
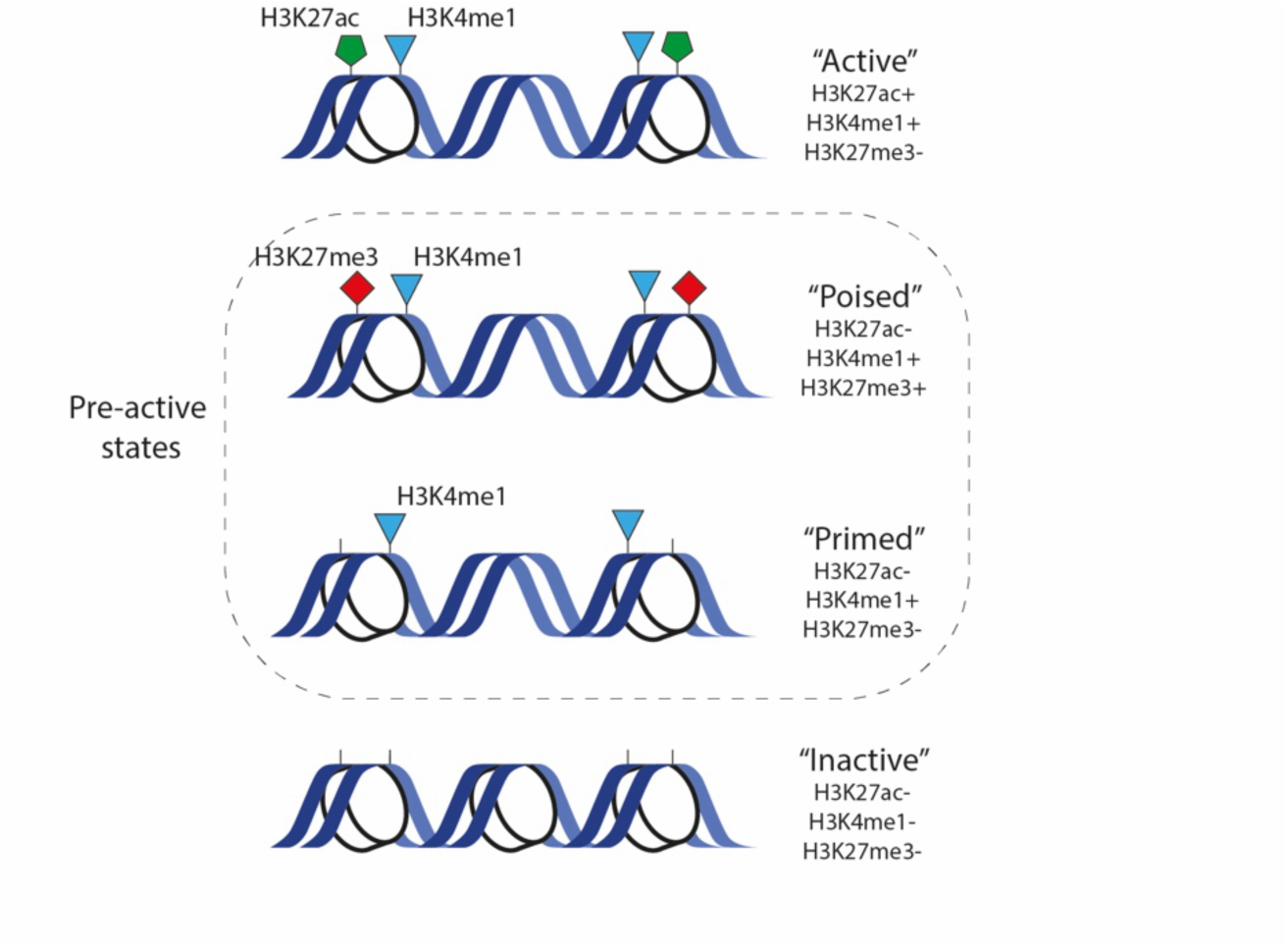
Schematics of the histone modification signatures associated with various enhancer activity states: Active (H3K27ac+/H3K4me1+/H3K27me3-), Poised (H3K27ac-/H3K4me1+/H3K27me3+), Primed (H3K27ac-/H3K4me1+/H3K27me3-), or Inactive (H3K27ac-/H3K4me1-/H3K27me3-).

**Supp. Figure 2.**
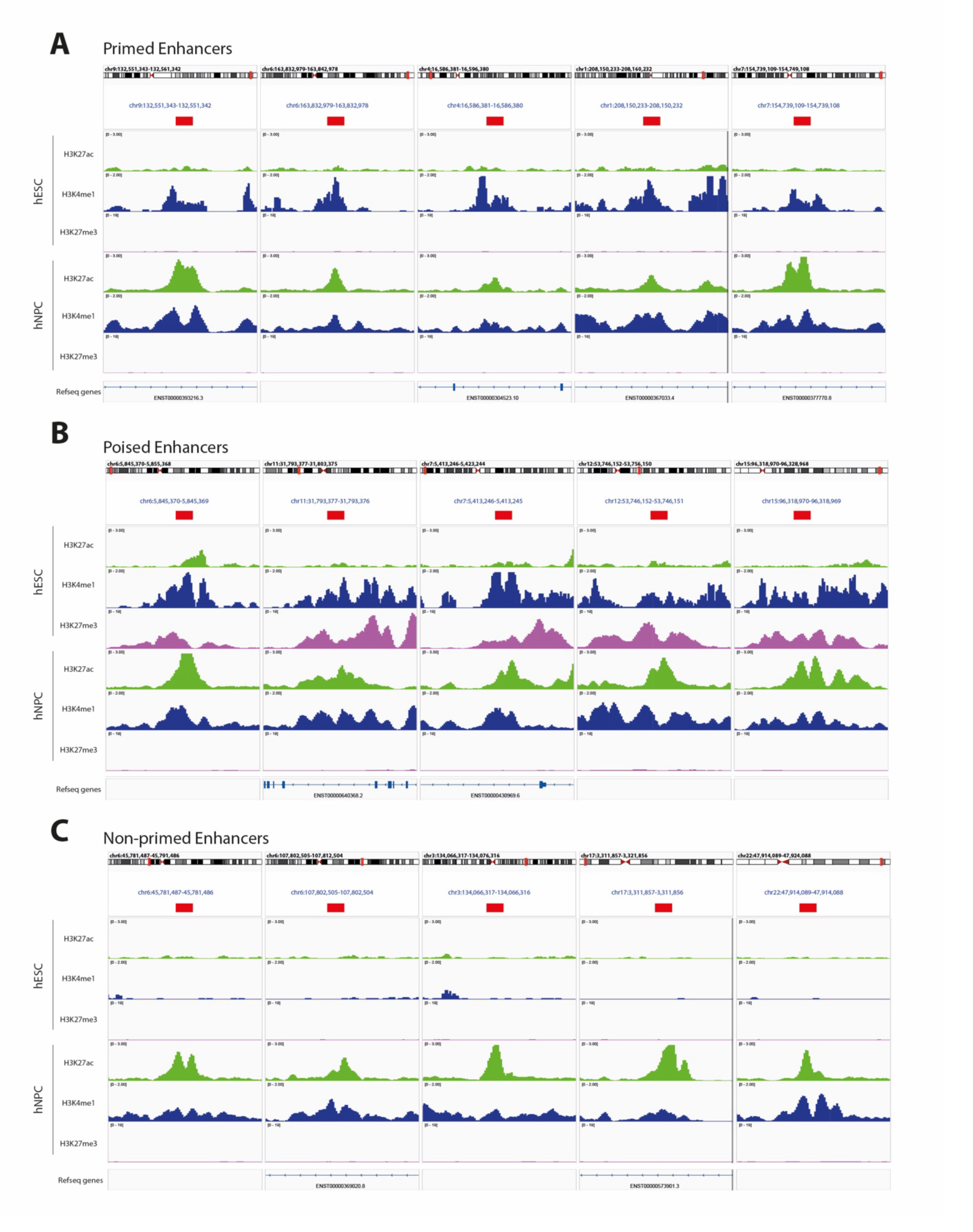
Genome browser visualization examples of hNPC enhancers that were identified as being either Primed **(A)**, Poised **(B),** or Non-primed **(C)** within hESCs. Each window represents a single enhancer instance with the enhancer annotated in red.

**Supp. Figure 3.**
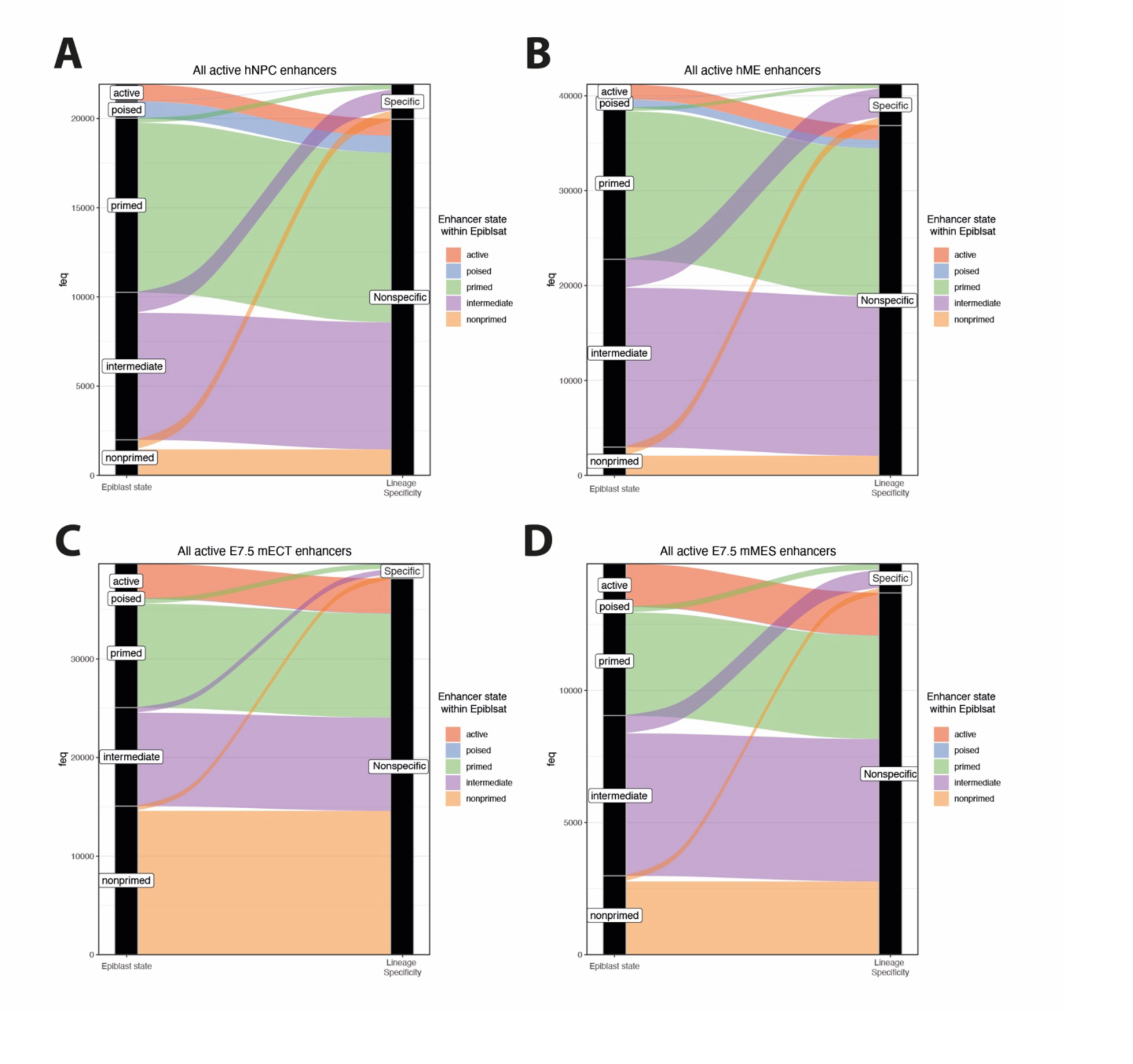
Alluvial plots of relative proportions of lineage-specific enhancers (H3K27ac signal in a single cell lineage) and nonspecific (H3K27ac signal at enhancer within multiple cell lineages) and their respective epigenetic state within hESC/mESC cultures. Plots for all identified hNPC enhancers **(A)**, hME enhancers **(B)**, E7.5 mECT enhancers **(C)**, and E7.5 mMES enhancers **(D)** are shown.

**Supp. Figure 4.**
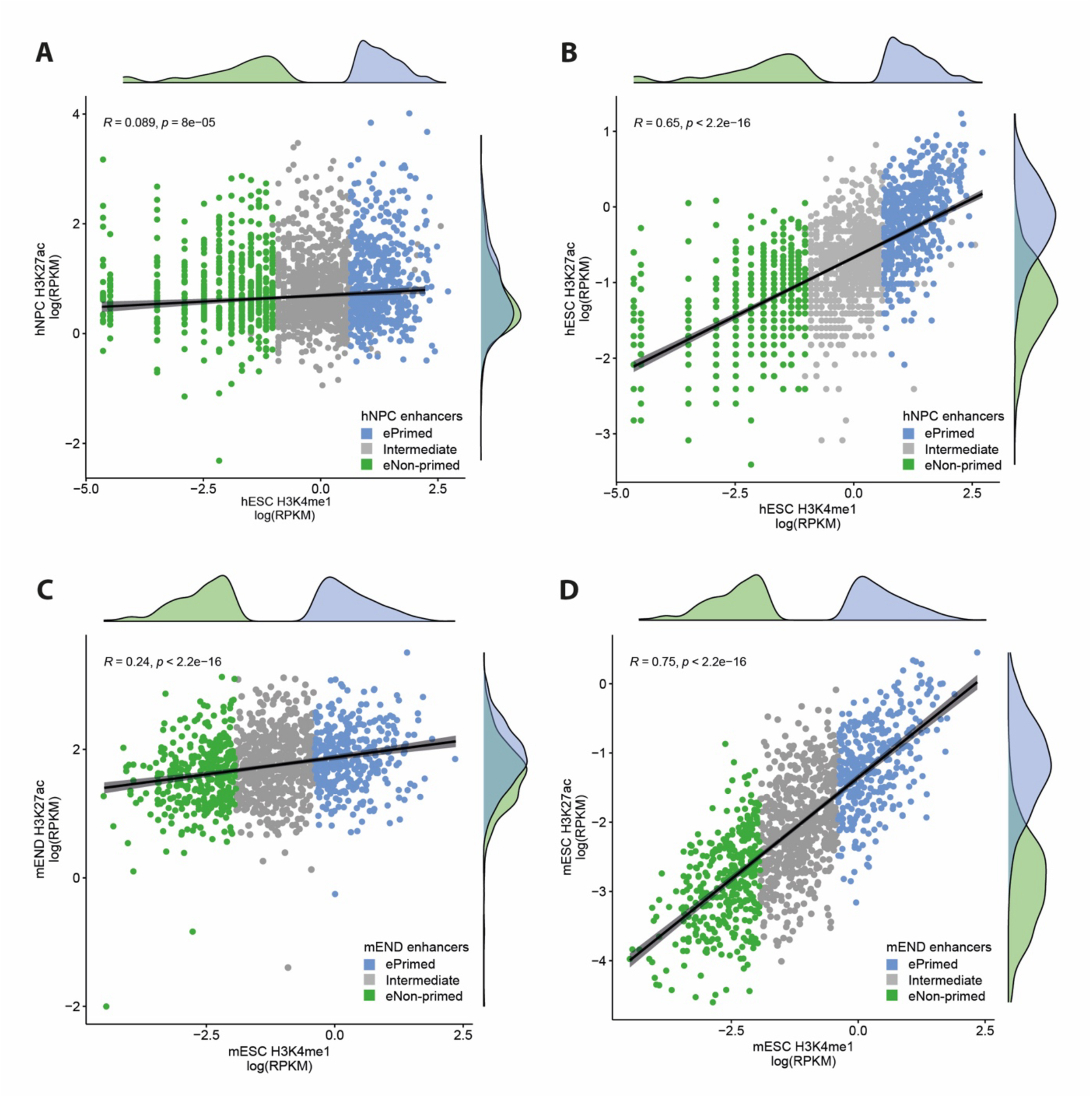
**(A-D)** ChIP-seq counts of lineage-specific primed (blue), non-primed (green) and intermediate (grey) sub-groups. Each dot is one enhancer element with a read count shown as log(RPKM) of 1.5kb window centred on the enhancer. Regression line of all lineage-specific enhancers within the group is shown with 95% confidence interval. Results of Pearson correlation coefficient test are also shown. Density plots of distribution of primed (blue) and non-primed (green) populations are shown on the x and y axes. **(A)** Read counts of hESC H3K4me1 and hNPC H3K27ac at hNPC enhancers. **(B)** Read counts of hESC H3K4me1 and hESC H3K27ac at hNPC enhancers. **(C)** Read counts of mESC H3K4me1 and mEND H3K27ac at E7.5 mEND enhancers. **(D)** Read counts of mESC H3K4me1 and mESC H3K27ac at E7.5 mEND enhancers.

**Supp. Figure 5.**
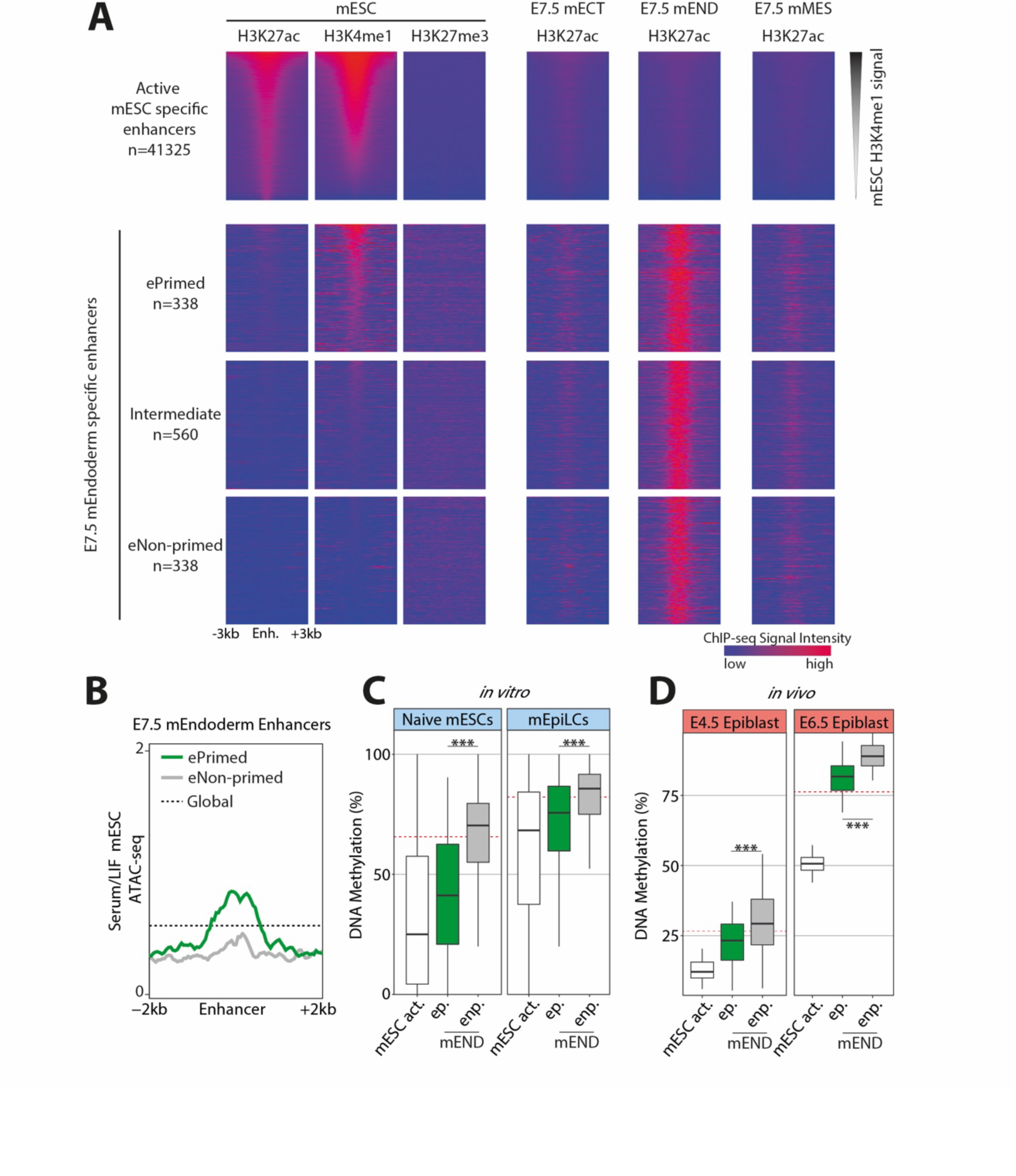
**(A)** Heatmaps of ChIP-seq H3K27ac, H3K4me1 and H3K27me3 data within mESCs, and of H3K27ac within *in vivo* dissections of mECT, mEND, and mMES tissues for E7.5 embryos. Shown here are enhancer elements with lineage-specific active states in mESC cells or E7.5 mEND tissue. Enhancer groups called as ePrimed (high H3K4me1, low H3K27ac), eNon-primed (low H3K4me1, low H3K27ac), or Intermediate (intermediate H3K4me1) are displayed. **(B)** ATAC-seq profile for E7.5 mEND called enhancer groups within *in vitro* mESCs. Running averages in 50-bp windows around the centre of the enhancer (2 kb upstream and downstream) are shown alongside the averaged global signal (black dashed line). **(C-D)** DNA methylation levels over 500bp core of active mESC enhancers (white) and mEND called enhancer sub-groups: ePrimed (ep.) and eNon-primed (enp.) **(C)** by WGBS within *in vitro* naive mESC/mEpiLCs and **(D)** by scNMT-seq of *in vivo* mouse embryos at days E4.5 and E6.5. **(C-D)** Box plots show median levels and the first and third quartile, whiskers show 1.5x the interquartile range. Global methylation levels are displayed (red dashed line) along with results of ANOVA test between primed and non-primed groups (p<0.001 ***).

**Supp. Figure 6.**
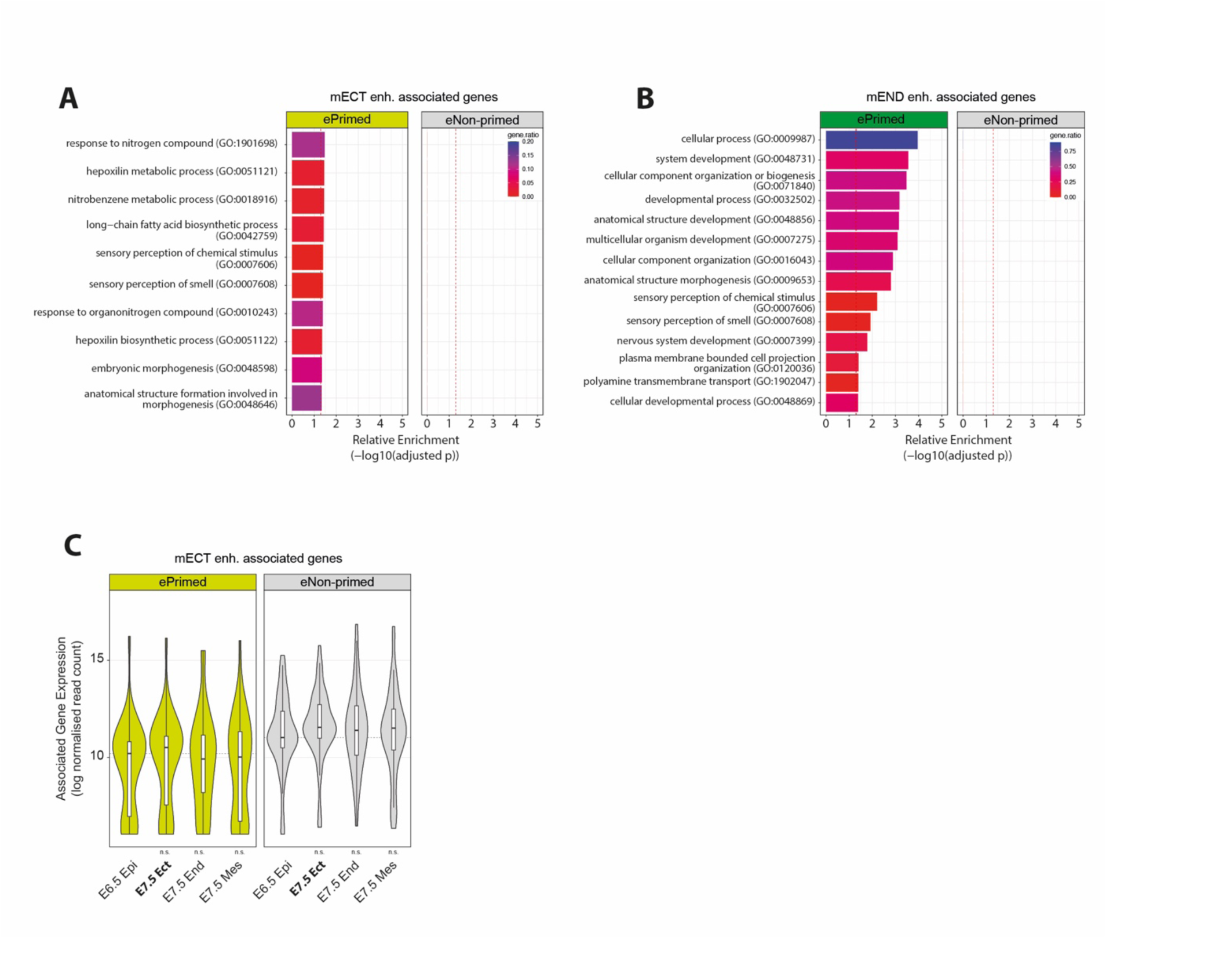
**(A-B)** Gene ontology enrichment analysis for **(A)** ePrimed and eNon-primed E7.5 mECT enhancer-associated genes, and **(B)** ePrimed and eNon-primed E7.5 mEND enhancer-associated genes. Significance threshold of p<0.05 is displayed (red dashed line). **(C)** Overlaid box and violin plots show expression of enhancer-associated genes. Box plots show median levels and the first and third quartile, whiskers show 1.5× the interquartile range. Results from Welch two-sample t-test against E6.5 epiblast are displayed. Expression (log normalized counts) within *in vivo* scNMT-seq data collected from E6.5 epiblast and E7.5 ectoderm/ endoderm/ mesoderm tissues of genes exclusively associated with either ePrimed E7.5 mECT (yellow) enhancers, or eNon-primed E7.5 mECT (grey) enhancers. Median expression for each gene group within E6.5 epiblast is also shown (grey dashed line).

**Supp. Figure 7.**
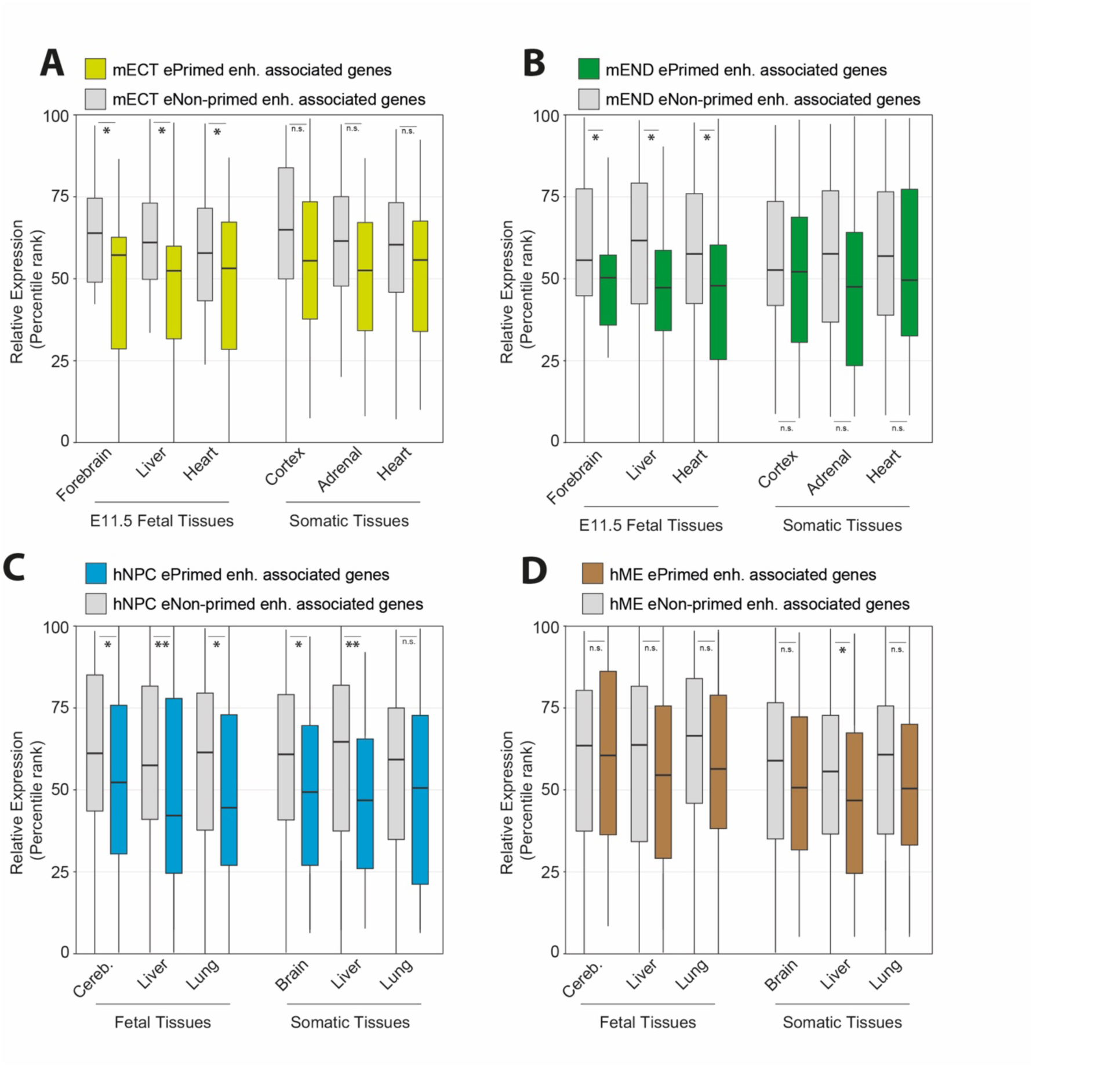
**(A-D)** Relative expression of enhancer-associated genes as defined by PCHiC/proximity hybrid model. Box plots show median levels and the first and third quartile, whiskers show 1.5x the interquartile range. Results from Wilcoxon signed-rank test between primed and non-primed enhancer sub-groups are displayed (p<0.01 **, p<0.05 *). **(A-B)** Relative expression within mouse E11.5 fetal and somatic tissues (2 month old), of mECT enhancer-associated genes **(A)** and of mEND enhancer-associated genes **(B)**. **(C-D)** Relative expression within human fetal tissues (72-129 post conception days) and somatic tissues (50-70 years of age), of hNPC enhancer-associated genes **(C)** and of hME enhancer-associated genes **(D)**.

**Supp. Figure 8.**
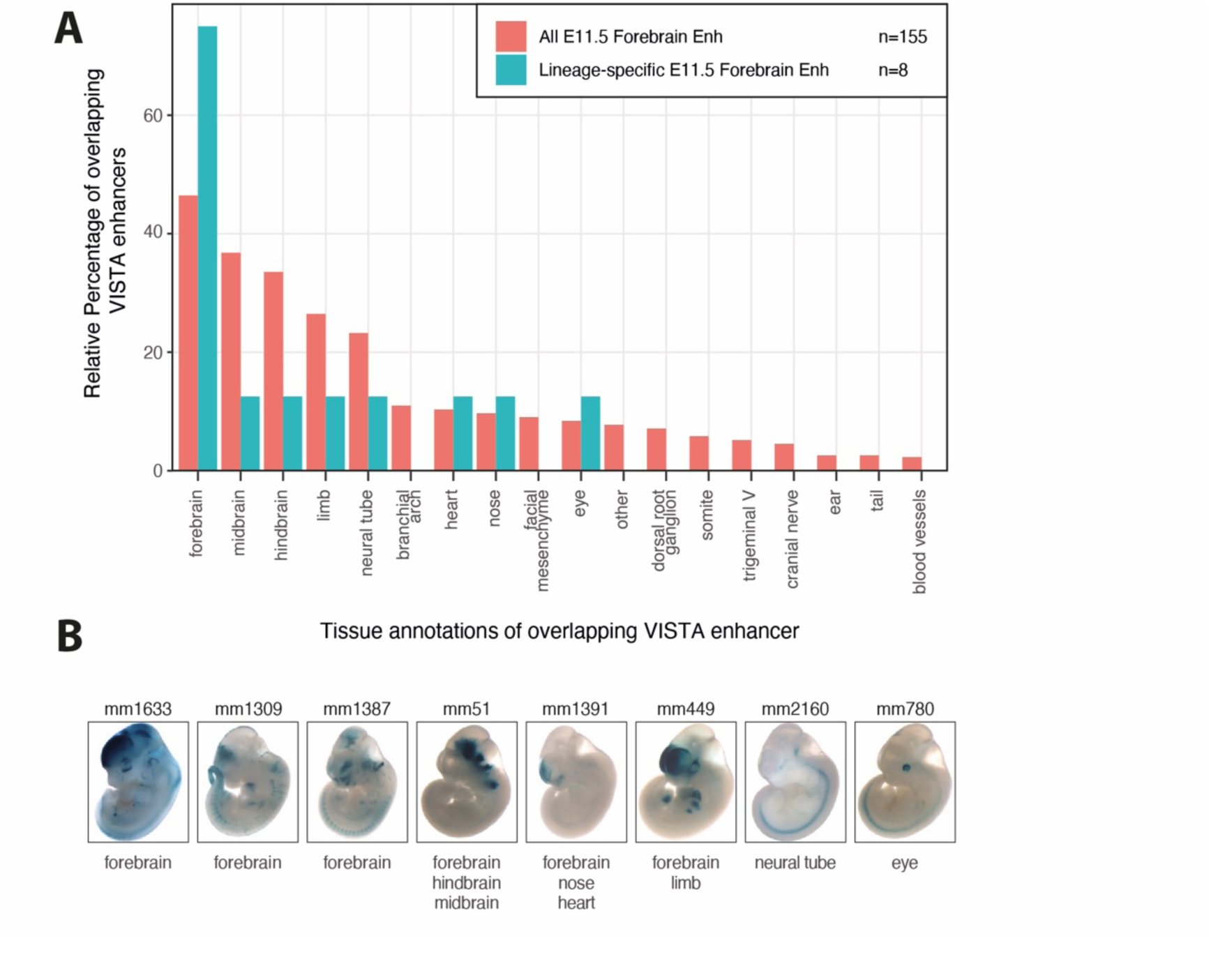
**(A)** Relative percentage of VISTA browser enhancers overlapping called E11.5 forebrain enhancers which are annotated for activity within respective embryonic tissues. VISTA enhancers overlapping all E11.5 forebrain enhancers are annotated in red, VISTA enhancers overlapping lineage-specific E11.5 forebrain enhancers are annotated in blue. **(B)** Representative images of reporter assay performed within E11.5 embryos for VISTA browser enhancers overlapping lineage-specific E11.5 forebrain enhancers. Tissues annotated with enhancer activity are listed below.

**Supp. Figure 9.**
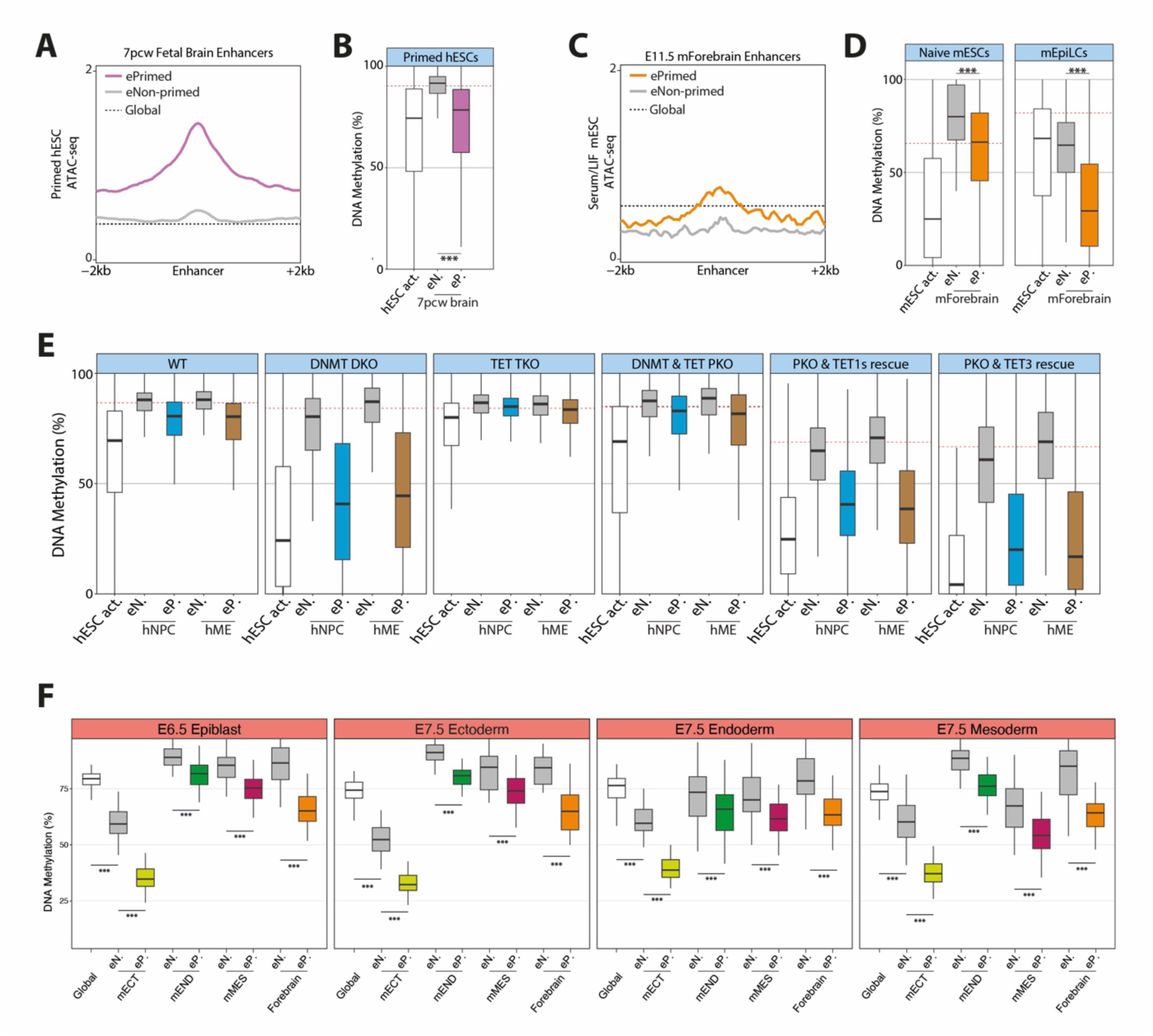
**(A)** ATAC-seq profile for 7pcw fetal brain called enhancer groups within *in vitro* hESCs. Running averages in 50-bp windows around the centre of the enhancer (2 kb upstream and downstream) are shown alongside the averaged global signal (black dashed line). **(B)** DNA methylation levels by WGBS within *in vitro* primed hESC over 500bp core of active hESC enhancers (white) and 7pcw fetal brain called enhancer sub-groups: ePrimed (ep.) and eNon-primed (enp.) **(C)** ATAC-seq profile for E11.5 forebrain called enhancer groups within *in vitro* mESCs. Running averages in 50-bp windows around the centre of the enhancer (2 kb upstream and downstream) are shown alongside the averaged global signal (black dashed line). **(D)** DNA methylation levels by WGBS within *in vitro* mESC over 500bp core of active mESC enhancers (white) and E11.5 forebrain called enhancer sub-groups: ePrimed (ep.) and eNon-primed (enp.). Global DNA methylation shown by red dashed line. **(E)** DNA methylation levels by WGBS within *in vitro* primed hESC with combinations of knockouts performed for DNMT and TET genes over 500bp core of active hESC enhancers (white) and hNPC and hME called enhancer sub-groups: ePrimed (ep.) and eNon-primed (enp.). Global DNA methylation shown by red dashed line. Knockout lines shown are wild type control (WT), DNMT3A/3B double knockout (DNMT DKO), TET1-3 triple knockout (TET TKO), DNMT3A/3B and TET1-3 pentuple knockout (DNMT & TET PKO), PKO line with TET1 rescue, and PKO line with TET3 rescue. **(F)** DNA methylation levels by scNMT-seq of *in vivo* mouse embryos at days E6.5 and E7.5 over 500bp core of E7.5 mECT, E7.5 mMES, E7.5 mEND, and E11.5 forebrain called enhancer sub-groups: eNon-primed (enp.) and ePrimed (ep.). Global methylation of all cells within the group are shown in white. **(B,D-F)** Box plots show median levels and the first and third quartile, whiskers show 1.5x the interquartile range. Results of ANOVA test between indicated groups are shown (p<0.001 ***).

**Supp. Figure 10.**
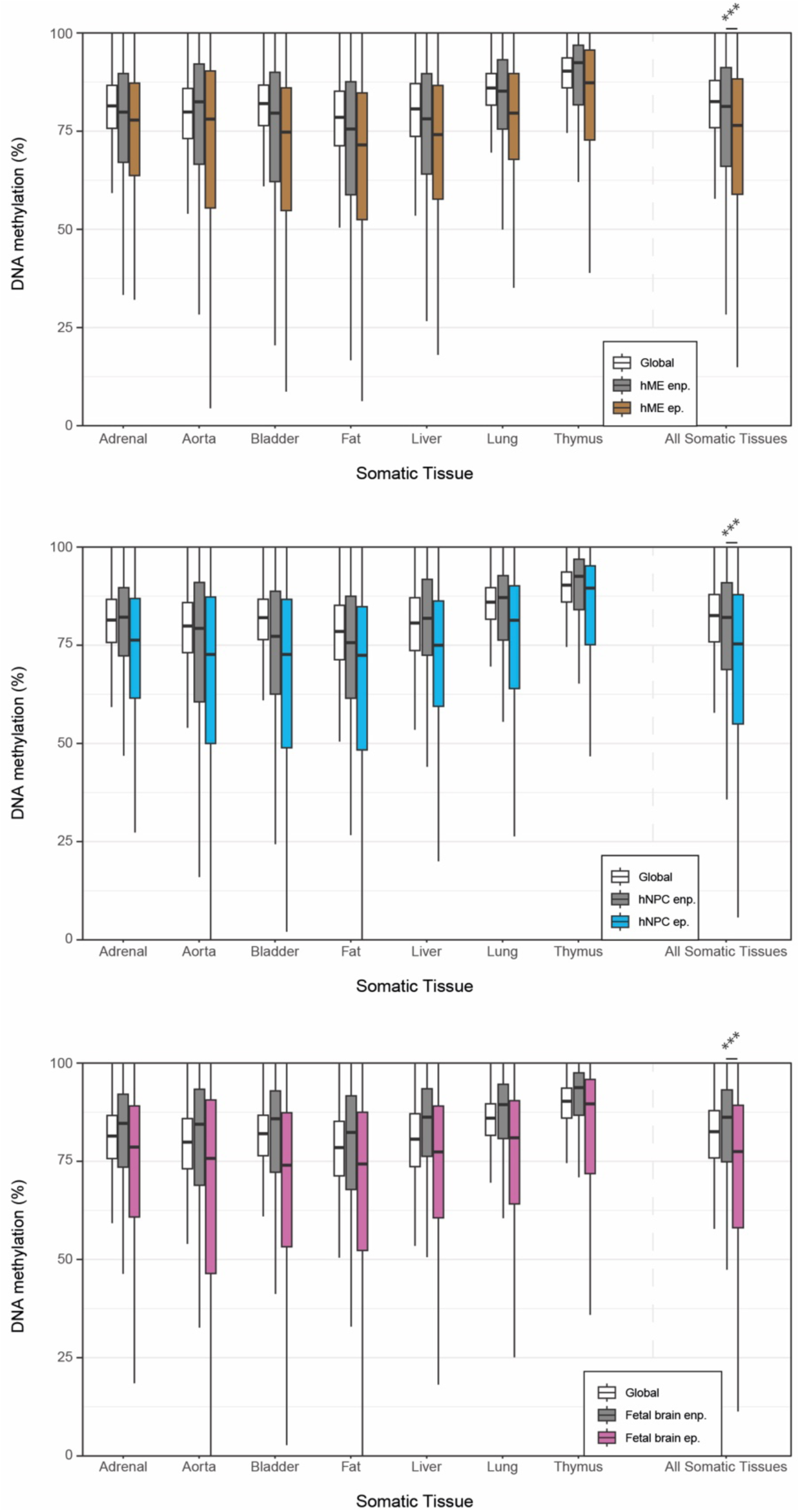
DNA methylation levels by WGBS-seq of *in vivo* human somatic tissues (50-70 years of age) over 500bp core of hME, hNPC and 7pcw fetal brain called enhancer sub-groups: eNon-primed (enp.) and ePrimed (ep.). Average methylation across all somatic tissues for each enhancer is also plotted. Box plots show median levels and the first and third quartile, whiskers show 1.5x the interquartile range. Global methylation calculated at 10kb running windows (white) are shown along with results of ANOVA test between indicated groups (p<0.001 ***).

**Supp. Figure 11.**
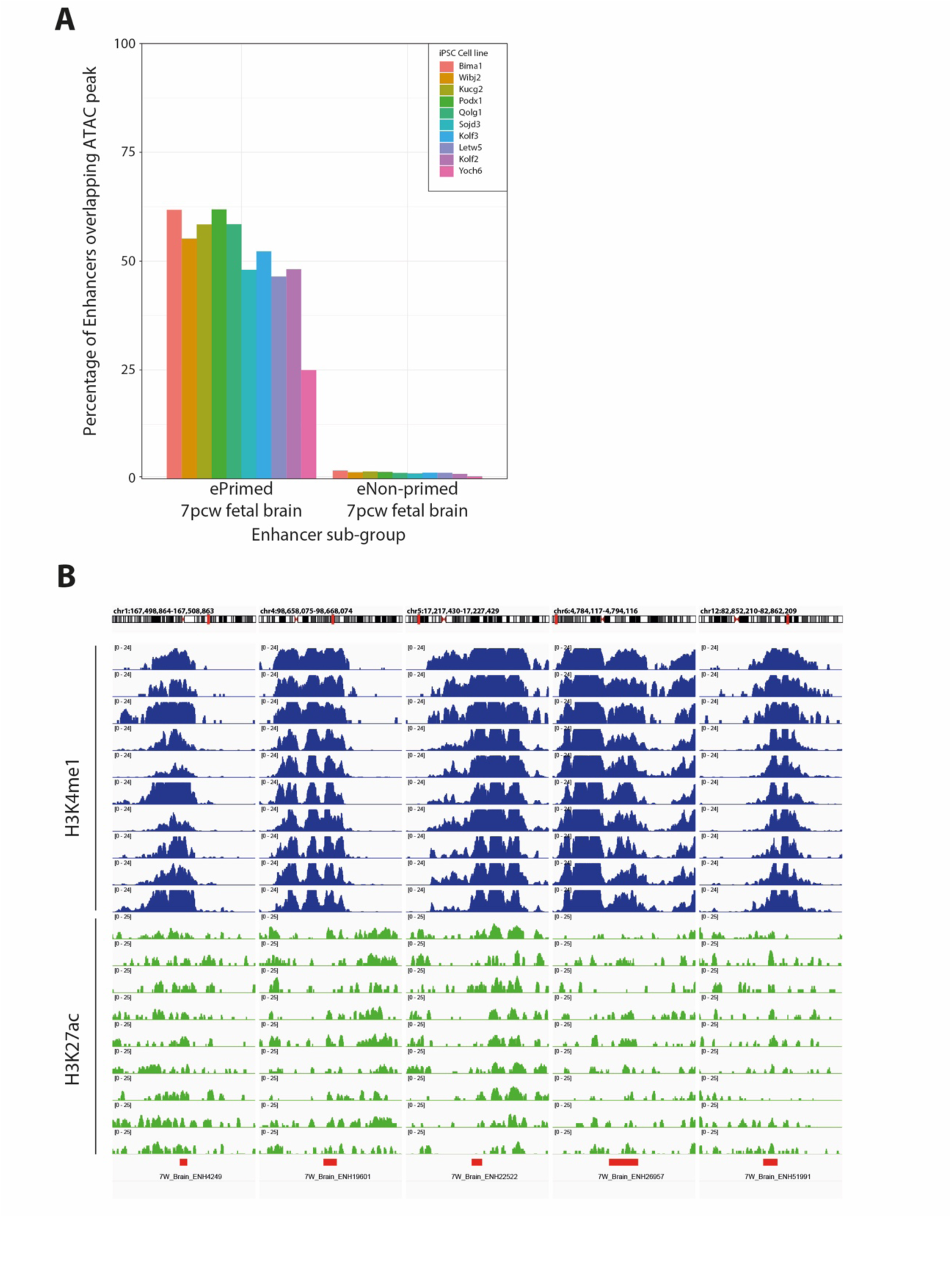
**(A)** Percentage of ePrimed and eNon-primed 7pcw fetal brain enhancers that overlap ATAC peaks within at least 2 of the 3 replicates per HipSci donor line. **(B)** Genome browser visualisation of respective examples of ePrimed 7pcw enhancers within the HipSci donor lines. Cut&Tag signal for H3K4me1 (blue) and H3K27ac (green) are shown. The centre for each enhancer is annotated below (red).

